# Western diet consumption impairs memory function via dysregulated hippocampus acetylcholine signaling

**DOI:** 10.1101/2023.07.21.550120

**Authors:** Anna M. R. Hayes, Logan Tierno Lauer, Alicia E. Kao, Shan Sun, Molly E. Klug, Linda Tsan, Jessica J. Rea, Keshav S. Subramanian, Cindy Gu, Natalie Tanios, Arun Ahuja, Kristen N. Donohue, Léa Décarie-Spain, Anthony A. Fodor, Scott E. Kanoski

**Affiliations:** Human and Evolutionary Biology Section, Department of Biological Sciences, University of Southern California, Los Angeles, CA, USA; Department of Bioinformatics and Genomics, University of North Carolina at Charlotte, Charlotte, NC, USA; Neuroscience Graduate Program, University of Southern California, Los Angeles, CA, USA

## Abstract

Western diet (WD) consumption during development yields long-lasting memory impairments, yet the underlying neurobiological mechanisms remain elusive. Here we developed an early life WD rodent model to evaluate whether dysregulated hippocampus (HPC) acetylcholine (ACh) signaling, a pathology associated with memory impairment in human dementia, is causally-related to WD-induced cognitive impairment. Rats received a cafeteria-style WD (access to various high-fat/high-sugar foods; CAF) or healthy chow (CTL) during the juvenile and adolescent periods (postnatal days 26-56). Behavioral, metabolic, and microbiome assessments were performed both before and after a 30-day healthy diet intervention beginning at early adulthood. Results revealed CAF-induced HPC-dependent contextual episodic memory impairments that persisted despite healthy diet intervention, whereas CAF was not associated with long-term changes in body weight, body composition, glucose tolerance, anxiety-like behavior, or gut microbiome. HPC immunoblot analyses after the healthy diet intervention identified reduced levels of vesicular ACh transporter in CAF vs. CTL rats, indicative of chronically reduced HPC ACh tone. To determine whether these changes were functionally related to memory impairments, we evaluated temporal HPC ACh binding via ACh-sensing fluorescent reporter *in vivo* fiber photometry during memory testing, as well as whether the memory impairments could be rescued pharmacologically. Results revealed dynamic HPC ACh binding during object-contextual novelty recognition was highly predictive of memory performance and was disrupted in CAF vs. CTL rats. Further, HPC alpha-7 nicotinic receptor agonist infusion during consolidation rescued memory deficits in CAF rats. Overall, these findings identify dysregulated HPC ACh signaling as a mechanism underlying early life WD-associated memory impairments.

## Introduction

Consumption of a Western diet (WD), broadly defined as a diet high in processed foods, saturated fat, and simple sugars, is associated with excessive caloric intake, obesity, and metabolic dysfunction (*1–4*). Independent of these outcomes, WD consumption is also linked with cognitive dysfunction (*5–7*), especially when consumed during early life periods of development (*8–10*). One region of the brain that is particularly vulnerable to early life dietary insults is the hippocampus (HPC) (*11, 12*), known for its key role in mediating episodic memory of previous experiences and the context (e.g., time, place) in which they occurred (*13–15*). An accumulating number of studies in both humans and animal models reveal that habitual WD consumption during early life leads to impairments in HPC-dependent learning and memory function, even absent of metabolic dysfunction or body weight gain (*16–20*). However, despite these established connections between WD consumption and disrupted memory, the underlying neurobiological mechanisms through which WD during development leads to long-lasting hippocampal dysfunction remain elusive.

WD- and obesity-associated alterations in HPC neural processes have been identified, including reduced levels of brain-derived neurotrophic factor, altered synaptic plasticity, and elevated markers of neuroinflammation (*21–25*). However, very little is understood about the underlying neuromodulators driving these outcomes. The HPC relies on acetylcholine (ACh) neurotransmission, particularly from the medial septum, for proper memory function (*26–29*) and disrupted ACh signaling is a pathological marker of Alzheimer’s disease (AD). There is a close relationship between amyloid β peptide (Aβ) accumulation and cholinergic dysfunction in AD, and Aβ suppresses the synthesis and release of ACh from septal neurons (*30–32*). Given the longitudinal associations between WD consumption and AD onset (*33*), disturbances in ACh signaling could be a mechanism for long-term WD-related memory impairments. Here we evaluate the long-lasting impact of early life WD consumption on HPC-dependent episodic memory, and the extent that behavioral outcomes are mediated by dysregulated HPC ACh signaling.

Dietary factors drastically alter the gut microbiome (*16, 34–38*) and a growing body of evidence supports a functional link between early life diet, cognitive function, and changes in gut bacteria (*7, 16, 39*). Based on this recent literature, including findings that WD-induced microbiome changes are associated with changes in brain acetylcholine (ACh) signaling (*40*), a plausible hypothesis is that the gut microbiome is functionally connected with early life WD-induced memory impairments, and potentially via changes in HPC ACh function. This hypothesis is evaluated here using an ethologically-relevant WD model that includes dietary choice (between various high sugar and/or fat food and drink options) and macronutrient profiles modeling a modern human WD.

The extent that WD-induced memory impairments are reversible with dietary intervention is poorly understood. Here, we aimed to elucidate how consumption of a WD in early life perturbs memory function in both the short- and long-term. Using our ‘junk food’ cafeteria-style diet model to represent an early life WD model in rats, we examined metabolic, behavioral, microbial, biochemical, and functional imaging outcomes after a 30-day WD access period from juvenile onset, as well as after a 30-day healthy diet intervention period starting in early adulthood. Our collective findings show that consumption of a WD in early life leads to long-lasting deficits in HPC-dependent episodic memory that are attributable to impaired HPC ACh signaling, which persist despite a healthy diet intervention in adulthood.

## Results

### Early life WD leads to minimal metabolic impairments

No differences in body weight were observed between CAF and CTL rats throughout either the WD access period or healthy diet intervention (Fig. 1A-B, Fig. S1A,C,E). Furthermore, no differences in body composition (lean/fat mass ratio) were observed either after the early life WD access period or after the healthy diet intervention period (Fig. 1C-D). Glucose tolerance was not altered in CAF rats compared to CTL rats at either time point (Fig. 1E-F). Despite the lack of body weight and body composition effects, rats receiving the CAF diet in early life consumed approximately 15% more kilocalories than CTL rats during the early life WD access period; however, this increased caloric intake did not persist when the CAF rats were switched to the healthy diet in adulthood (Fig. 1G; Fig. S1B,D,F). During the WD access period, rats in the CAF group consumed the majority of their kilocalories from the distinct CAF components in the following order (highest to lowest % kcal consumed): high-fat, high-sugar chow > peanut butter cups ≥ potato chips > HFCS (Fig. 1H; Supplementary Fig. S2A-C). In terms of macronutrients during the WD period, CAF rats consumed approximately equal kilocalories (44%) from fat and carbohydrate, and the remaining from protein (12%) (Fig. 1I; Fig. S2A-C). For comparison, the healthy standard chow that the CTL group received throughout the study and that the CAF group received for the healthy diet intervention contained 13%, 58%, and 29% kcal from fat, carbohydrate, and protein, respectively. Collectively, these results show that despite promoting overconsumption of energy, the WD model did not induce an obesogenic weight trajectory or metabolic impairments, and the hyperphagia reversed immediately with the healthy diet intervention.

**Fig. 1.**
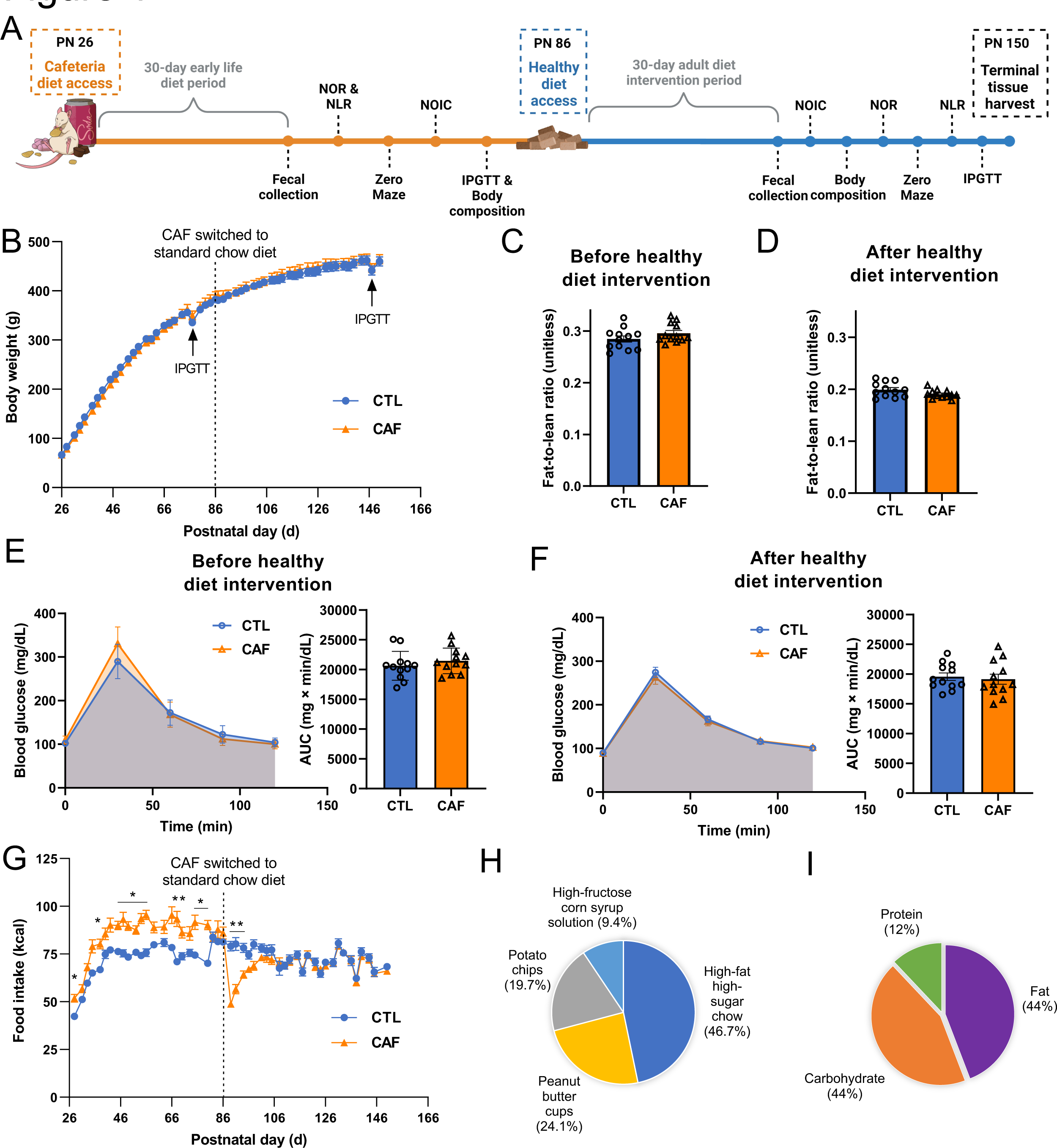
Early life Western diet consumption does not result in immediate or persistent alterations in body weight trajectory, body composition, or glucose tolerance, despite promoting hyperphagia during the Western diet (WD) period. (**A**) Experimental design timeline for behavioral and metabolic assessments. (**B**) Body weight over time (postnatal day) throughout the course of study for the main metabolic and behavioral cohort (N=24 total rats, n=12 CTL, n=12 CAF; two-way ANOVA with repeated measures over time; time × diet: P=0.0004, no post hoc differences using Sidak’s multiple comparisons test, time: P<0.0001, diet: P=0.8772; additional cohorts are shown in Fig. S1). (**C**) Assessment of body composition (fat-to-lean ratio) following the WD period but before the healthy diet intervention (N=24 total rats, n=12 CTL, n=12 CAF; unpaired t-test [2-tailed]; P=0.0766; additional body composition values can be found in Fig. S2). (**D**) Assessment of body composition (fat-to-lean ratio) following the healthy diet intervention (N=24 total rats, n=12 CTL, n=12 CAF; unpaired t-test [2-tailed]; P=0.1834; additional body composition values can be found in Fig. S2). (**E**) Glucose tolerance test results after the WD period but before the healthy diet intervention (N=24 total rats, n=12 CTL, n=12 CAF; two-way ANOVA (diet, time [as repeated measure], diet × time interaction) for blood glucose over time; unpaired t-test [2-tailed] for blood glucose AUC; time × diet: P<0.0001, no post hoc differences using Sidak’s multiple comparisons test, time: P<0.0001, diet: P=0.3812, for AUC: P=0.3861). (**F**) Glucose tolerance test results following the healthy diet intervention (N=24 total rats, n=12 CTL, n=12 CAF; two-way ANOVA (diet, time [as repeated measure], diet × time interaction) for blood glucose over time; unpaired t-test [2-tailed] for blood glucose AUC; time × diet: P=0.8630, time: P<0.0001, diet: P=0.7111, for AUC: P=0.6909). (**G**) Food intake expressed as kcal over time (postnatal day) throughout the course of study for the main metabolic and behavioral cohort (N=24 total rats, n=12 CTL, n=12 CAF; two-way ANOVA with repeated measures over time; time × diet: P<0.0001, many post hoc differences using Sidak’s multiple comparisons test, time: P<0.0001, diet: P=0.0479; additional cohorts are shown in Fig. S1). (**H**) Percentage of energy intake consumed from the CAF diet components for the main metabolic and behavioral cohort (N=24 total rats, n=12 CTL, n=12 CAF; additional cohorts are shown in Fig. S1). (I) Percentage of energy intake from macronutrients for the main metabolic and behavioral cohort (N=24 total rats, n=12 CTL, n=12 CAF; additional cohorts are shown in Fig. S2). AUC, area under the curve; CAF, cafeteria diet group; CTL, control group; WD, Western diet. Error bars represent ± SEM. *P<0.05, **P<0.01.

### Early life WD imparts long-lasting HPC-dependent memory impairments without influencing perirhinal cortex-dependent memory or novelty aversion

We performed both Novel Location Recognition (NLR) and Novel Object in Context (NOIC) to assess HPC-dependent spatial recognition (*41, 42*) and contextual episodic memory (*43, 44*), respectively (Fig. 2A-F). Following the early life WD access period, there were no differences in spatial recognition memory evaluated through NLR (Fig. 2B); however, after the healthy diet intervention period the CAF rats showed an impairment in this task (Fig. 2C). As for NOIC, deficiencies in contextual episodic memory were apparent both before and after the healthy diet intervention period in CAF rats (Fig. 2E-F; Fig. S3), signifying that early life WD consumption led to long-lasting memory impairments, regardless of consumption of a healthier diet in adulthood. Importantly, these deficits occurred absent of any differences in total object exploration time between the CAF and CTL groups (Fig. S3A-D,F-G,H-K). Together, these findings reveal that early life WD ‘programs’ long-lasting deficits in HPC-dependent memory function.

**Fig. 2.**
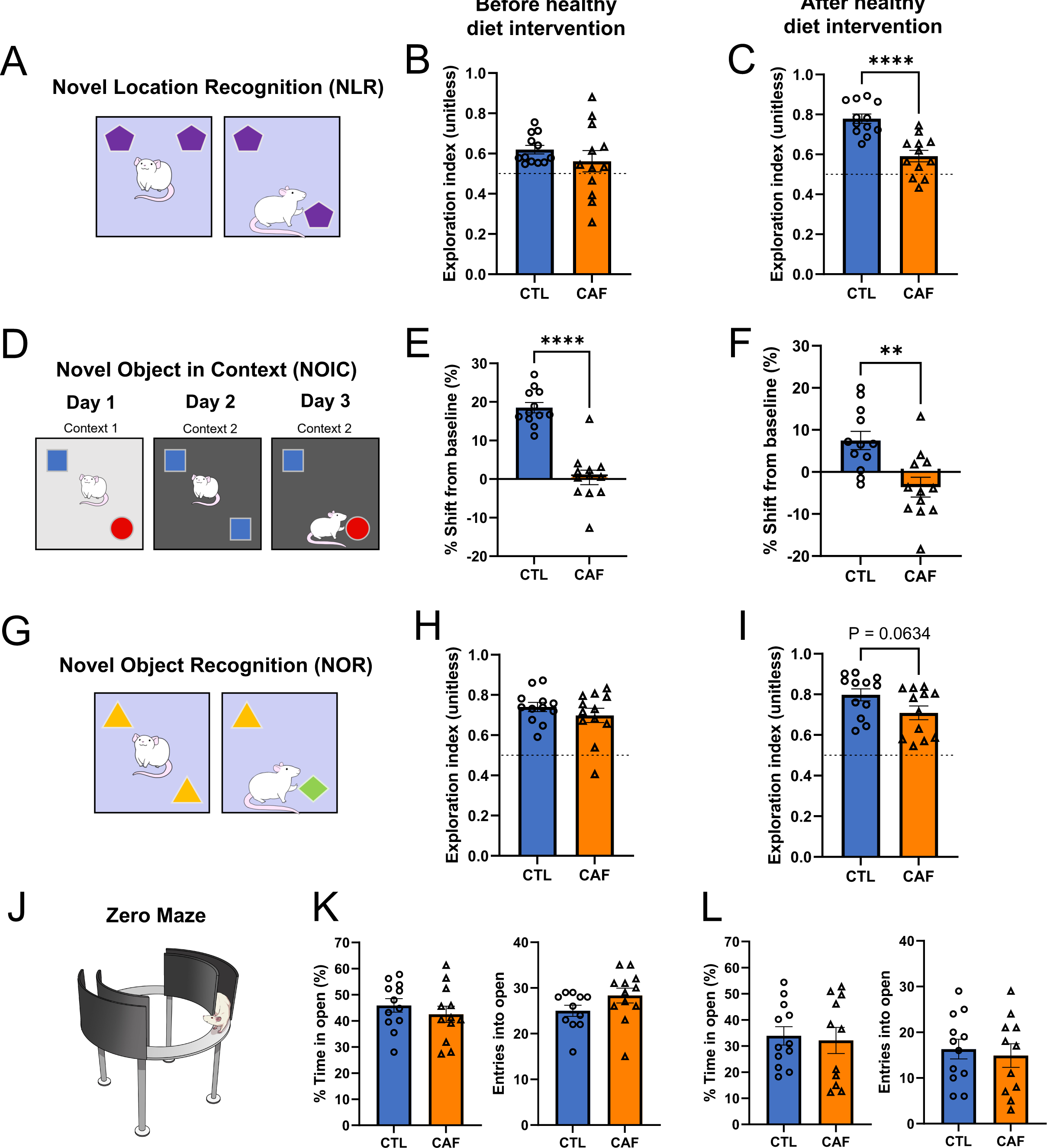
Early life Western diet (WD) consumption yields long-lasting episodic memory impairments in the absence of impacts on perirhinal cortex-dependent novel object exploration or markers of anxiety. (**A**) Novel Location Recognition (NLR) behavioral scheme. (**B**) NLR exploration index following the WD access period but before the healthy diet intervention (N=24 total, n=12 CTL, n=12 CAF; unpaired t-test [2-tailed] with Welch’s correction for unequal variances; P=0.3258). (**C**) NLR exploration index following the healthy diet intervention (N=24 total, n=12 CTL, n=12 CAF; unpaired t-test [2-tailed]; P<0.0001). (**D**) Novel Object in Context (NOIC) behavioral scheme. (**E**) NOIC percent shift from baseline performance following the WD access period but before the healthy diet intervention (N=24 total, n=12 CTL, n=12 CAF; unpaired t-test [2-tailed]; P<0.0001). (**F**) NOIC percent shift from baseline performance following the healthy diet intervention (N=24 total, n=12 CTL, n=12 CAF; unpaired t-test [2-tailed]; P=0.002). (**G**) Novel Object Recognition (NOR) behavioral scheme. (**H**) NOR exploration index following the WD access period but before the healthy diet intervention (N=24 total, n=12 CTL, n=12 CAF; unpaired t-test [2-tailed]; P=0.3320). (**I**) NOR exploration index following the healthy diet intervention (N=24 total, n=12 CTL, n=12 CAF; unpaired t-test [2-tailed]; P=0.0634). (**J**) Zero Maze apparatus illustration. (**K**) Percent time spent in and entries into the open arm regions of the Zero Maze apparatus after the WD access period but before the healthy diet intervention (N=23 total, n=11 CTL due to one outlier, n=12 CAF; unpaired t-test [2-tailed]; P=0.4064 for percent time in open, P=0.1175 for entries into open). (**L**) Percent time spent in and entries into the open arm regions of the Zero Maze apparatus following the healthy diet intervention (N=23 total, n=12 CTL, n=11 CAF due to one outlier; unpaired t-test [2-tailed]; P=0. 7719 for percent time in open, P=0. 6752 for entries into open). CAF, cafeteria diet group; CTL, control group; WD, Western diet. Error bars represent ± SEM. **P<0.01, ****P<0.0001.

To evaluate object recognition memory that is not HPC-dependent, we determined whether consumption of a WD in early life impairs memory in a Novel Object Recognition (NOR) behavioral approach that relies on the perirhinal cortex independently of the HPC (*45, 46*) (Fig. 2G). Immediately following the early life WD access period, there were no differences in exploration index of a novel vs. non-novel object between CAF and CTL rats (Fig. 2H; P=0.33). Following the healthy diet intervention period, there was a trend for decreased exploration of a novel object in CAF rats vs. CTL rats, but this did not reach significance (Fig. 2I; P=0.06). These findings indicate early life WD consumption did not lead to robust differences in recognition of a novel object in a memory task that does not involve the HPC. Importantly, these results suggest that NLR and NOIC impairments were not secondary to novelty aversion.

### Early life WD does not alter markers of anxiety or locomotor activity

Because memory and anxiety-like behavior can both involve hippocampal function (*47–49*), we examined markers of anxiety-like behavior and locomotor activity through the Zero Maze and Open Field tests. There were no differences in the percentage of time that rats spent in the open arms of the Zero Maze apparatus, a marker of anxiety-like behavior, between the CAF and CTL groups either before or after the healthy diet intervention (Fig. 2J-L).

Furthermore, there were no differences in the number of entries into the open arms of the Zero Maze apparatus at either time point (Fig. 2K-L), which is an assessment of locomotor activity. We also performed the Open Field task after the WD access period, and there were no differences in distance travelled, an additional assessment of locomotor activity, or time spent in the center of the apparatus, an additional marker of anxiety-like behavior, in this test (Fig. S3L-M). These results signify that the persistent HPC-dependent memory impairments due to a WD are not confounded by altered anxiety-like behavior or locomotor activity.

### Early life WD leads to gut microbiome dysbiosis that is reversed with adult healthy diet intervention

Because the gut microbiome has previously been shown to be altered by WD exposure (*35*), we examined whether early life WD consumption impacted gut microbial taxonomic composition both immediately after the WD access period as well as after the healthy diet intervention period. Cladogram visualization and PCoA of 16s rRNA sequencing of fecal samples collected immediately after the 30-day WD period reveal robust differences in gut microbiome at the genus level between CAF and CTL rats (Fig. 3A-B). Similar differences were observed in the PCoA for sequences of cecal content of animals that were not exposed to the healthy diet intervention (Fig. 3D). However, after the healthy diet intervention, PCoAs of sequences of both fecal and cecal contents revealed that the initial marked differences were largely reversed (Fig. 3C,E). Shannon indices, which assess α-diversity of microbial communities in fecal and cecal samples were significantly lower in the CAF group than those in the CTL group after the early life WD access period, but these differences did not persist after the healthy diet intervention (Fig. S4). A cladogram for the time point after WD exposure depicts the broad distribution of taxa across groups (Fig. 3A). By contrast, only 5 taxa were significantly altered after the healthy diet intervention (FDR<0.1), further indicating that early life WD did not impart long-lasting effects on the gut microbiome.

**Fig. 3.**
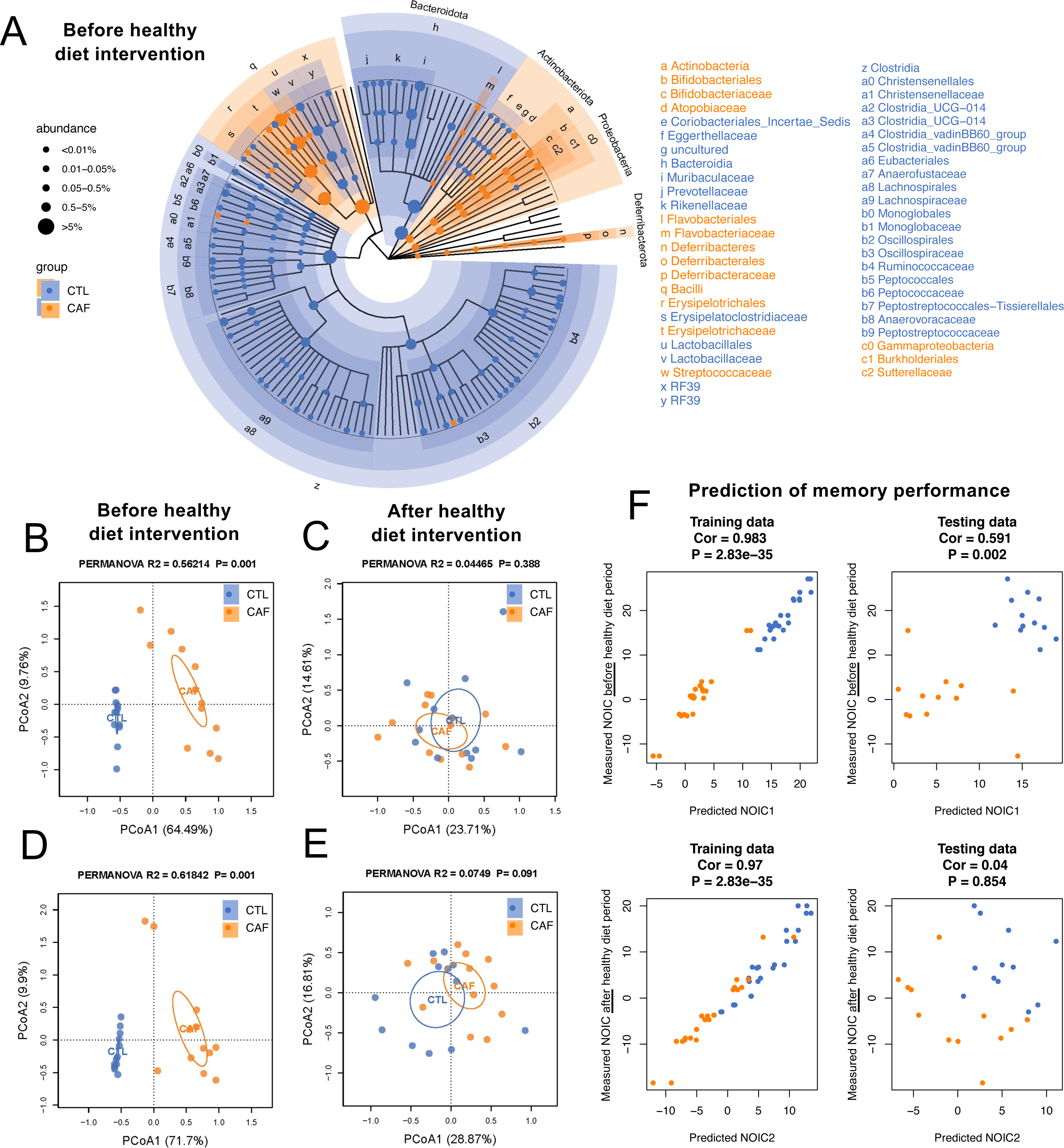
Early life Western diet (WD) robustly alters the gut microbiome, but these alterations do not persist upon a healthy diet intervention in adulthood. (**A**) Cladogram representation of differentially abundant taxa in fecal matter after the WD period but before the healthy diet intervention period (N=24 total, n=12 CTL, n=12 CAF; false discovery rate [FDR] <0.1 for significant taxa). (**B**) Principal coordinate analysis (PCoA) of microbiomes at the genus level in fecal matter after the WD access period (N=24 total, n=12 CTL, n=12 CAF; Bray-Curtis dissimilarity; PERMANOVA, R^2^=0.56214, P=0.001, P<0.05 for significant associations). (**C**) PCoA of microbiomes at the genus level in fecal matter after the healthy diet intervention period (N=24 total, n=12 CTL, n=12 CAF; Bray-Curtis dissimilarity; PERMANOVA, R^2^=0.04465, P=0.388). (**D**) PCoA of microbiomes at the genus level in cecal content after the WD access period (N=24 total, n=12 CTL, n=12 CAF; Bray-Curtis dissimilarity; PERMANOVA, R^2^=0.61842, P=0.001). (**E**) PCoA of microbiomes at the genus level in cecal content after the healthy diet intervention period (N=24 total, n=12 CTL, n=12 CAF; Bray-Curtis dissimilarity; PERMANOVA, R^2^=0.0749, P=0.388). (**F**) Machine learning analysis to evaluate whether microbiome composition before the healthy diet intervention period reliably predicts NOIC memory performance either before or after the healthy diet period (N=24 total, n=12 CTL, n=12 CAF; random forest regression models, 3-fold cross validation; for training data: Cor 0.983 and P=2.83e-35 [before healthy diet intervention period], Cor 0.97 and P=2.83e-35 [after healthy diet intervention period]; for testing data: Cor 0.591 and P=0.002 [before healthy diet intervention period], Cor 0.04 and P=0.854 [after healthy diet intervention period]). CAF, cafeteria diet group; CTL, control group; FDR, false discovery rate; PCoA, principal coordinate analysis; WD, Western diet.

Given the pronounced differences in gut microbiome immediately after the early life WD access period, we performed correlation analyses to determine if any microbial taxa were related to the early life WD memory impairments observed. A number of correlational analyses between specific taxon and memory performance in the NOIC task at the time point following the early life WD access period withstood the FDR correction (Fig. S4). Most notably, individual abundances of the genera *Lactococcus* and *Bifidobacterium* were negatively correlated with NOIC memory performance (Fig. S4). In contrast, abundance of the species *Lactobacillus intestinalis* was positively correlated with NOIC memory performance (Fig. S4). We then sought to decipher a potential mechanistic role of the gut microbiome in the long-lasting memory deficits by building a machine learning model to test whether the gut microbiome immediately after the WD access period could predict memory function in NOIC after the healthy diet intervention. Using random forest regression models with both 3-fold and 4-fold cross validation, we observed that microbiome after the WD period was linked with NOIC memory performance before the healthy diet intervention but not later in life after the healthy diet intervention (Fig. 3F, Fig. S5). This indicates that the microbiome was not responsible for the long-lasting memory impairments observed.

### Early life WD yields long-lasting reductions in chronic HPC ACh tone

Given that the HPC relies on ACh neurotransmission for proper memory function (*26–29*), we examined levels of proteins related to ACh signaling in the HPC as a potential novel mechanism for enduring HPC dysfunction from early life WD. Protein quantification was evaluated for choline acetyltransferase (ChAT), an enzyme that synthesizes ACh from the precursors choline and acetyl-CoA whose expression correlates positively with spatial memory (*50*); vesicular ACh transporter (VAChT), which transports ACh to vesicles for synaptic release and whose expression is positively correlated with improved spatial memory function during aging (*51*); and HPC acetylcholinesterase (AChE), an enzyme that degrades ACh and that has been found to disrupt memory function (*52, 53*). Immunoblotting analyses from dorsal HPC tissue collected after the healthy diet intervention period revealed that there were no differences in ChAT or AChE levels, yet CAF rats had reduced levels of VAChT compared to CTL rats (Fig. 4A-C). These results suggest that there was no post-synaptic compensation for a reduced amount of ACh reaching a synapse. To further determine whether these changes in chronic cholinergic tone could be related to features of the microbiome, we performed correlational analyses between the abundances of key microbial taxa both before and after the healthy diet intervention and VAChT levels, and results revealed no significant correlations for any taxa (Fig. S6). Overall, these findings suggest that early life WD consumption leads to long-lasting reductions in cholinergic tone.

**Fig. 4.**
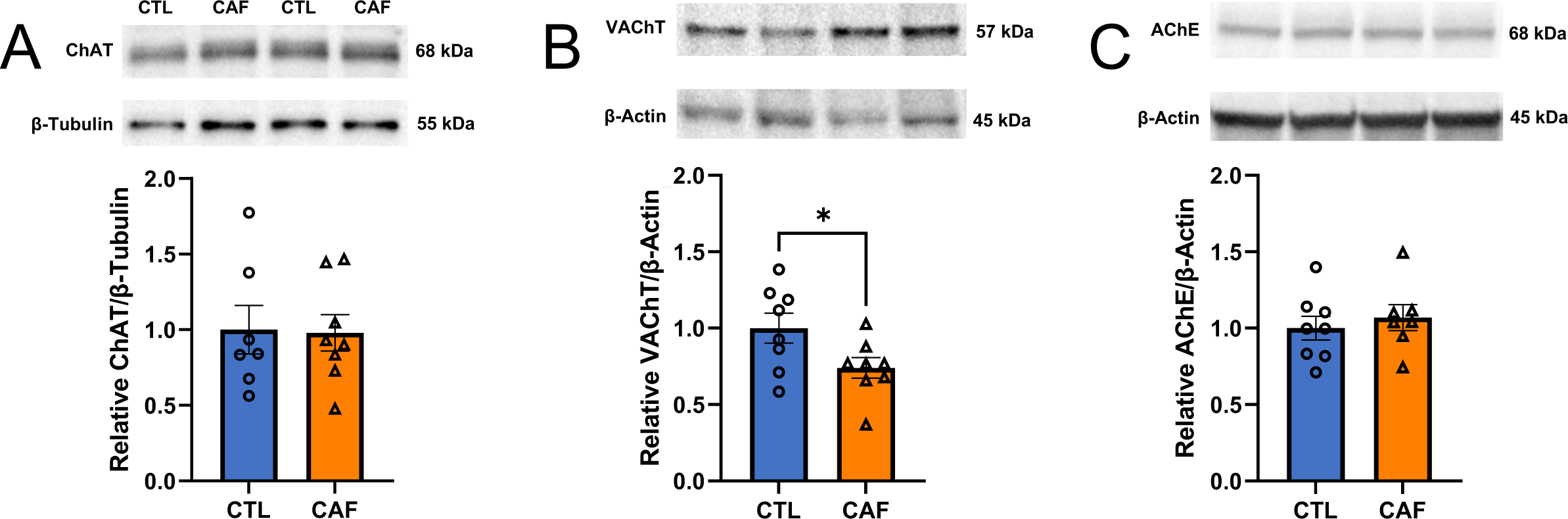
Early life Western diet (WD) leads to long-lasting reduction in hippocampal cholinergic tone. (**A**) Immunoblot representative images and results for choline acetyltransferase (ChAT) protein levels in the dorsal hippocampus (HPC) of rats following WD access in early life and a healthy diet intervention in adulthood (N=15 total, n=7 CTL due to outlier, n=8 CAF; unpaired t-test [2-tailed], P=0.9165). (**B**) Immunoblot representative images and results for vesicular acetylcholine transporter (VAChT) protein levels in the dorsal HPC of rats following early life WD access and a healthy diet intervention in adulthood (N=16 total, n=8 CTL, n=8 CAF; unpaired t-test [2-tailed], P=0. 0449). (**C**) Immunoblot representative images and results for acetylcholinesterase (AChE) protein levels in the dorsal HPC of rats following WD access in early life and a healthy diet intervention in adulthood (N=15 total, n=8 CTL, n=7 CAF due to outlier; unpaired t-test [2-tailed], P=0. 5570). ACh, acetylcholine; AChE, acetylcholinesterase; CAF, cafeteria diet group; ChAT, choline acetyltransferase; CTL, control group; HPC, hippocampus; VAChT, vesicular acetylcholine transporter; WD, Western diet. Error bars represent ± SEM. *P<0.05.

### Early life WD disrupts acute ACh signaling dynamics during memory testing

Given that early life WD consumption resulted in enduring memory dysfunction and decreases in HPC ACh tone, we next investigated whether acute ACh signaling dynamics during the contextual episodic memory task were altered in CAF vs. CTL rats. After the 30-day healthy diet intervention period, CAF and CTL animals underwent the NOIC behavioral task with simultaneous recording of ACh signaling via *in vivo* fiber photometry (Fig. 5A-B). Replicating the results from our previous cohorts, results revealed that CAF rats were impaired in the task (Fig. 5C). CTL rats showed increased ACh binding at the moment of investigating the object novel to the context compared to the object familiar to the context on the test day (day 3; Fig. 5D, Fig. S7), whereas CAF rats showed no difference in ACh binding upon investigating the two objects on the test day (Fig. 5E, Fig. S7). Comparing between groups, the extent of ACh binding at the onset of exploring the object novel to the context was significantly elevated in CTL rats compared to CAF rats (Fig. 5F), but there were no differences in ACh binding between groups at the onset of investigating the object familiar to the context (Fig. 5G). Additionally, there were no differences in ACh binding when investigating the objects upon initial exposure to them on NOIC day 1 (Fig. S7). Collectively, these findings reveal early life WD access has lasting effects on the temporal dynamics of ACh signaling in the HPC during discrimination of a context-object novelty.

**Fig. 5.**
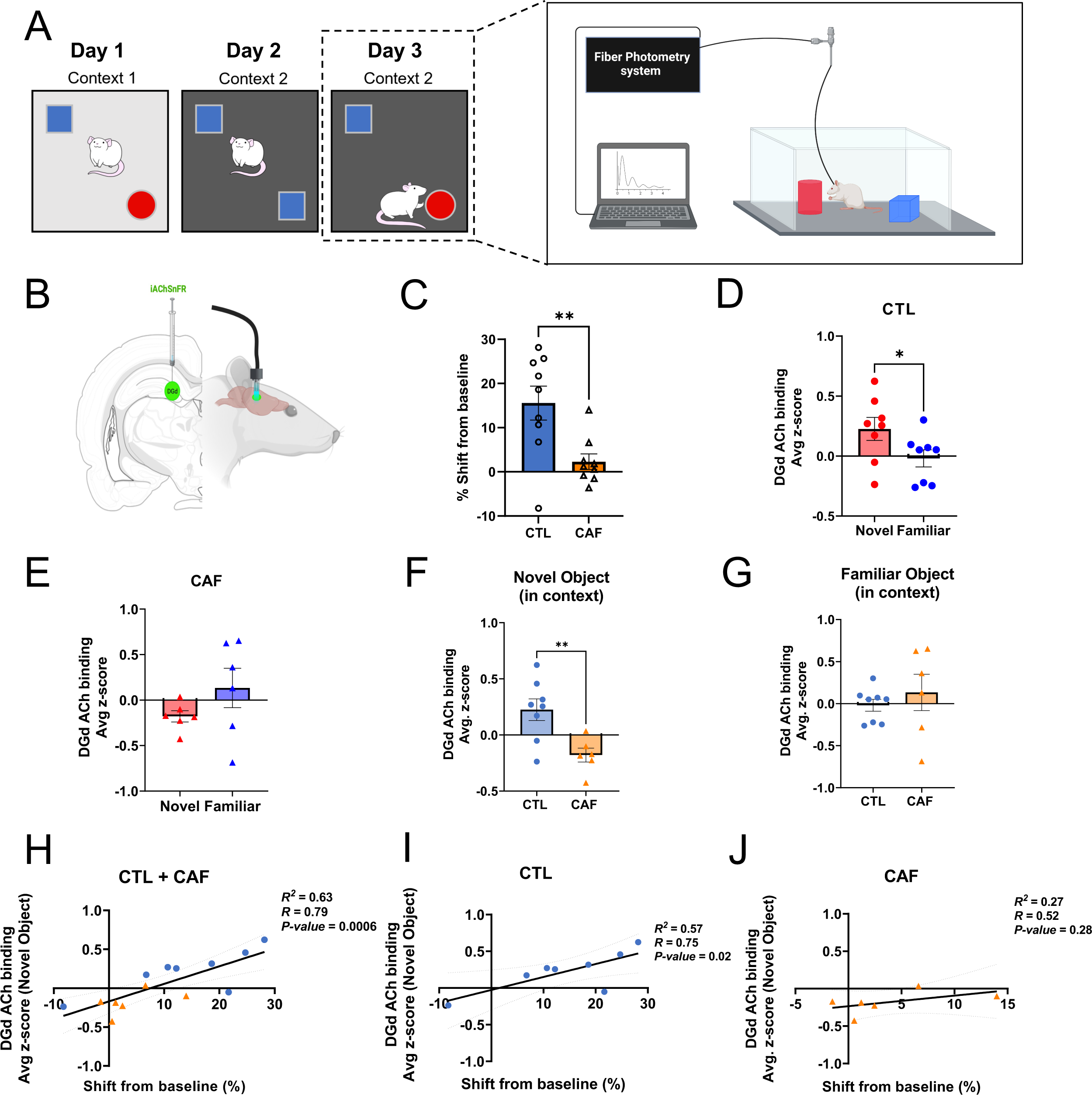
Early life Western diet (WD) disrupts hippocampal cholinergic binding upon contextual-object novelty encounter during episodic memory testing. (**A**) Behavioral scheme for the Novel Object in Context contextual episodic memory task coupled with *in vivo* fiber photometry. (**B**) In vivo fiber photometry approach to track acetylcholine (ACh) binding in the dentate gyrus (DG) region of the dorsal hippocampus (HPC). (**C**) NOIC percent shift from baseline performance following healthy diet intervention and with implementation of *in vivo* fiber photometry (N=18 total, n=9 CTL, n=9 CAF; unpaired t-test [2-tailed]; P=0.0064). (**D**) Average z-score of ACh binding in the DG at the onset of exploring the novel vs. familiar objects during the test day of NOIC in CTL rats (N=8 total, n=8 CTL group only, paired t-test [2-tailed], P=0.0140). (**E**) Average z-score of ACh binding in the DG at the onset of exploring the novel vs. familiar objects during the test day of NOIC in CAF rats (N=6 total, n=6 CAF group only, paired t-test [2-tailed], P=0. 1453). (**F**) Comparison of average z-score of ACh binding in the DG at the onset of exploring the novel object during the test day of NOIC in CTL vs. CAF rats (N=14 total, n=8 CTL, n=6 CAF; unpaired t-test [2-tailed], P=0.0068). (**G**) Comparison of average z-score of ACh binding in the DG at the onset of exploring the familiar object during the test day of NOIC in CTL vs. CAF rats (N=14 total, n=8 CTL, n=6 CAF; unpaired t-test [2-tailed], P=0. 4641). (**H**) Regression of average z-score of ACh binding at the onset of novel object exploration on percent shift from baseline NOIC memory performance (N=14 total, n=8 CTL, n=6 CAF; R^2^=0.63, P=0.0006). (**I**) Regression of average z-score of ACh binding at the onset of novel object exploration on percent shift from baseline NOIC memory performance in CTL rats only (N=8 total, n=8 CTL; R^2^=0.57, P=0.02). (**J**) Regression of average z-score of ACh binding at the onset of novel object exploration on percent shift from baseline NOIC memory performance in CAF rats only (N=6 total, n=6 CAF; R^2^=0.27, P=0.28). ACh, acetylcholine; CAF, cafeteria diet group; CTL, control group; DG, dentate gyrus; HPC, hippocampus; NOIC, Novel Object in Context; WD, Western diet. Error bars represent ± SEM. *P<0.05, **P<0.01.

To determine whether these ACh binding photometric data were related to memory performance, we regressed the extent of HPC DGd ACh binding with memory performance (shift from baseline in the NOIC task) with both CAF and CTL groups combined and observed a strong positive correlation, such that the better the rats did in the memory task, the greater the ACh binding during object-contextual novelty encounter (Fig. 5H). When separated by diet group, the positive correlation remained for the CTL group of rats (Fig. 5I) but dissipated for the CAF group (Fig. 5J). These results indicate that in CTL rats, with intact ACh neurotransmission, better memory performance was linked with increased ACh signaling upon object-context novelty encounter, whereas no such link was found in CAF rats with disrupted ACh neurotransmission.

### ACh agonists reverse early life WD-induced memory impairments

Given the long-term reduction in HPC ACh tone and altered temporal HPC ACh dynamics in CAF rats, we next sought to determine whether pharmacological administration of ACh receptor agonists could rescue the long-lasting deficits in memory function in CAF rats. On day 2 of the HPC-dependent NOIC task (tested after a healthy diet intervention), each rat was given bilateral infusions of either a general ACh receptor agonist (AChRa; carbachol), an α7 nicotinic receptor ACh agonist (α7nAChRa; PNU 282987), or vehicle 3-8 minutes immediately prior to undergoing the task (Fig. 6A-B). On the subsequent test day of the task, rats that received either AChRa (carbachol) or α7nAChRa (PNU) exhibited improved memory performance compared to rats receiving vehicle alone, with the latter group showing memory performance at chance levels analogous CAF rats in preceding experiments (Fig. 6C). These findings not only reveal that early life WD-induced memory impairments are reversed by augmenting ACh transmission during the consolidation phase of the memory task, but also that α7 nicotinic receptor signaling likely underlies the WD-induced memory dysfunction.

**Fig. 6.**
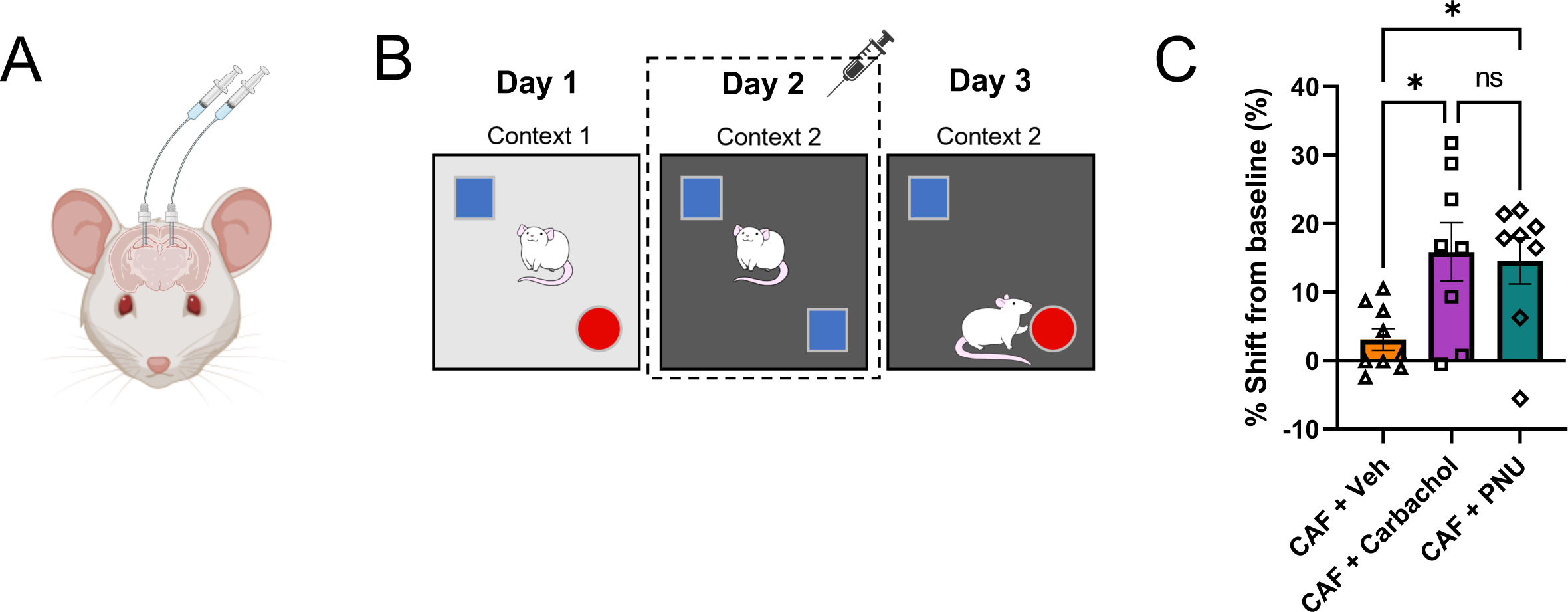
Pharmacological administration of acetylcholine agonists improves Western diet-induced memory performance. (**A**) Illustration of bilateral indwelling catheter infusions into the dorsal HPC. (**B**) Schematic of the Novel Object in Context (NOIC) contextual episodic memory task depicting timing of acetylcholine (ACh) agonist infusion on day 2. (**C**) NOIC percent shift from baseline performance following healthy diet intervention and with ACh infusion on day 2 (N=25 total, n=9 CAF+Vehicle, n=8 CAF+Carbachol [general ACh agonist], n=8 CAF+PNU [α7 nicotinic receptor agonist] due to outliers; One-way ANOVA (infusion group) with Tukey’s post-hoc test for multiple comparisons; P=0.0149 for overall model, for multiple comparisons: CAF+Vehicle vs. CAF+Carbachol: P=0.0226, CAF+Vehicle vs. CAF+PNU: P=0.0431, CAF+Carbachol vs. CAF+PNU: P=0.9542). ACh, acetylcholine; CAF, cafeteria diet group; CTL, control group; HPC, hippocampus; NOIC, Novel Object in Context; WD, Western diet. Error bars represent ± SEM. *P<0.05.

## Discussion

Although the negative metabolic impacts of consuming a Western diet (WD) high in refined sugar, saturated fat, and processed foods have been widely investigated (*1–4*), much less is known regarding the mechanisms through which WD leads to cognitive impairment. Here we developed an early life rodent WD model that, absent of metabolic or general behavioral abnormalities, yields enduring memory deficits that persist even after healthy diet intervention at the onset of adulthood. Results reveal that these long-lasting early life WD-induced memory impairments are mediated by compromised hippocampus (HPC) acetylcholine (ACh) signaling, manifested both in terms of chronic reductions in HPC ACh tone as well as aberrant acute temporal ACh binding dynamics in the HPC during mnemonic evaluation. The translational relevance of these findings is enhanced by additional results revealing that HPC administration of an ACh agonist targeting α7nAChRs during the memory consolidation phase reversed the WD-induced memory impairments, thus identifying potential pharmacological targets for diet-induced cognitive impairment.

While perturbations in the gut microbiome were observed during the early life WD-access period, these outcomes are unlikely tied to be to the memory deficits, as healthy diet intervention in adulthood reversed the microbiome changes but failed to rescue the memory impairments. Collectively, these findings denote a mechanistic role of ACh signaling, and not the gut microbiome, in persistent memory impairments due to WD consumption during early life periods of development.

The reliance of the HPC on ACh for proper memory function and the connection between impaired ACh neurotransmission and Alzheimer’s disease (AD) have been well-established (*26, 29–32*). The current findings identify altered ACh signaling as a functional mediator of early life WD-induced alterations in memory function. Previous evidence in adult rats receiving either a WD containing 22% kcal from fat, a high-fat diet containing 60% kcal from fat, or a control diet containing 7% kcal from fat for 12 weeks showed no differences in levels of ChAT in the medial septum/vertical limb of the diagonal band or in levels of AChE in the HPC despite impairments in spatial memory for the WD and high-fat diet groups (time spent in the novel arm of a Y-maze apparatus) (*54*). Integrated with our present results, this suggests early life (juvenile-adolescent periods) is a critical period for dietary exposure to impact HPC ACh neuronal pathway development. This is further supported by previous research indicating that administration of choline (an ACh precursor) throughout gestation and up through PN 24 enhanced spatial memory of rats in adulthood (*55*). In contrast, a recent study in diet-induced obese adult rats revealed increased levels of ChAT in the HPC after 5 and 17 weeks’ consumption of a high-fat diet containing 45% kcal from fat, decreased levels of AChE in the HPC after 17-week consumption of the diet, and increased level of VAChT in the HPC after 17-week consumption of the diet compared to control chow-fed rats (*56*). Furthermore, expression of the α7nAChR (nicotinic) was decreased at both 5 and 17 weeks on the diet in the HPC, while no effects were observed for muscarinic ACh receptor subtypes 1, 3, or 5 in the HPC regardless of time point (*56*). With the exception of the decreased level of α7nAChR, these findings suggest that ACh neurotransmission was increased in animals fed a high-fat diet, yet an important factor for consideration is that no behavior tests were performed to assess HPC function or other cognitive tasks. In addition, the study design differed in age of the rats and the duration of diet exposure compared to our present work. However, this previous study did identify that α7nAChR is more susceptible to high-fat dietary insults, which aligns with our current finding that selective agonism of α7nAChR rescued early life WD-induced HPC memory impairments.

Given this connection between diet, memory, and ACh signaling, an imperative topic of future research is to specifically determine how diet is ultimately giving rise to altered ACh neurotransmission and, furthermore, whether this leads to a predisposition to develop AD or other forms of dementia. We hypothesize a role for gastrointestinal (GI)-originating vagus nerve signaling in mediating WD-associated alterations in HPC ACh signaling. For example, we recently demonstrated that ablation of GI-specific vagal sensory signaling impairs HPC-dependent contextual episodic memory in rats (*57*). In this report we also identified the medial septum as an anatomical relay connecting vagal signaling to the dorsal HPC. Given that the medial septum is a principal source of ACh input to the HPC, it may be that blunted GI-derived vagal signaling contributes to the altered ACh signaling associated with early life WD. While additional work is required to support this hypothesis, it is the case that WD consumption is associated with blunted GI vagal signaling (*58–60*), thus providing additional evidence for this working model (*57*).

Our results clearly show that consumption of WD in early life markedly altered the gut microbiome, but, when the animals were switched to a healthy diet in adulthood, the alterations were largely reversed. The initial changes in the gut microbiome between CAF and CTL rats was expected, considering that the diets for each group were very distinct with marked differences in dietary fiber content, dietary fiber type (e.g., source, structure), macronutrient composition, micronutrient characteristics, and energy density, among other factors, which have been linked with shifts in gut microbial taxonomic populations in both humans and animal studies (*37, 61–63*). However, persistence of diet-induced taxonomic shifts in the gut microbiome have varied by age of exposure to a given diet, duration of diet exposure, type of diet used, host genome, and sex, among a variety of other potential factors (*64, 65*). In our previous work with the same early life CAF diet model implemented in females, gut microbiome differences following the healthy diet intervention became more disparate (*35*) instead of being reversed as with the current study in males. Previous research has shown that male and female mice have distinctly different bacterial taxonomic compositions as well as bacterial diversity in adulthood, independent of diet (*66, 67*). Given that machine learning analyses in the present study indicate that early WD-associated microbiome changes were not associated with long- (after dietary intervention) term memory impairments in males, and that males and females display divergent WD-induced gut microbiome phenotypes in adulthood yet both have long-lasting HPC-dependent cognitive deficits (*35*), we conclude that the gut microbiome is unlikely to be mediating the enduring HPC dysfunction associated with early life WD. To further support this and lend evidence that the microbiome is not related to altered HPC ACh neurotransmission, our correlational analyses between microbial taxa and HPC VAChT levels in male rats revealed no significant relationships. Regardless, whether aberrant HPC ACh signaling is functionally connected to WD-associated memory impairments in females requires further experimentation to fully elucidate.

An important distinction of the current findings is that the persistent WD-induced memory impairments due to altered ACh signaling occurred in the absence of effects on metabolic outcomes and body weight. This signifies that early life diet can have a critical, long-lasting effects on neural function – independently of obesity. Previous research has also found HPC-dependent memory impairments in the absence of differences in body weight or metabolic markers following early life free-access to 11% sugar or 11% HFCS in rats (*68*), or before the onset of body weight differences in mice or rats fed a high-fat diet (*69, 70*).

These findings suggest that development of HPC-dependent neurocognitive deficits may precede weight gain and obesity phenotype. However, in the present study the CAF rats were hyperphagic compared to controls during the CAF exposure period. One possibility for this caloric intake-body weight mismatch is based on a micronutrient deficiency in the CAF group. This is unlikely, however, as the predominant source of calories in the CAF model was the commercially-available high-fat diet, which provides sufficient micronutrient consumption. A more likely explanation is that the CAF model altered energy expenditure. However, putative changes in energy expenditure are unlikely to account for the HPC dysfunction given that there was no caloric intake-body weight mismatch when memory impairments were observed after the healthy dietary intervention.

The present study has some limitations to be considered. For example, the CAF diet model used does not allow for determination of which specific dietary component(s) or macronutrient(s) give rise to the effects observed. Further, we did not directly investigate a role for sex and/or sex hormones. Given that our previous work in female rats showed early life WD-access led to initial deviations in the gut microbiome that became more apparent after the healthy diet intervention (vs. controls) – an effect not observed in males – it is possible that early life WD exposure has sex-specific effects. As is inherent with all animal models, the translation to humans is nonlinear. However, it should be underscored that WD consumption is also linked with memory impairment AD in humans (*33, 71*). Given that AD is associated with aberrant HPC ACh signaling, the present results may provide important translational insight into links between the early life dietary environment and late-life neurocognitive impairments.

The present findings collectively reveal a mechanistic connection between early life exposure to a WD, long-lasting contextual episodic memory impairments, and ACh neurotransmission. The α7 nicotinic receptor was found to be a key intermediary of the WD-induced disrupted memory function, and persistent memory deficits were observed independent of altered metabolic outcomes or general behavior abnormalities – and were not specifically linked to dysbiosis of the gut microbiome. Overall, this work identifies a link between early life diet and impaired ACh neurotransmission in the HPC. Whether these findings have translational relevance to the etiology of human dementia warrants further investigation.

## Materials and Methods

### Subjects

The subjects for all experiments were male Sprague Dawley rats obtained from Envigo (Indianapolis, IN, USA) and housed in the animal vivarium at the University of Southern California. Rats arrived at the animal facility on postnatal day (PN) 25 and started their respective diet on PN 26. They were housed under a reverse 12:12 h light-dark cycle, with lights turned off at 11:00 and turned on at 23:00, and with temperature of 22-24°C and humidity of 40-50%. All rats were singly housed in hanging wire cages in order to facilitate food spillage collection for accurate measurement of individual food intake. Animals were weighed three times per week between 8:30-10:30 (shortly before the onset of the dark cycle). The specific numbers of animals used in each experiment are provided in the general experimental overview below as well as in Supplementary Table S1. All animal procedures were approved by the University of Southern California Institutional Animal Care and Use Committee (IACUC) in accordance with the National Research Council Guide for the Care and Use of Laboratory Animals.

### Dietary model

We implemented a junk food cafeteria-style diet (CAF) to model a Western diet in the present experiments, a model that we previously developed in female rats (*35*). This CAF diet consisted of free-choice *ad libitum* access to high-fat high-sugar chow (HFHS diet; Research Diets D12415, Research Diets, New Brunswick, NJ, USA; 20% kcal from protein, 35% kcal from carbohydrate, 45% kcal from fat), potato chips (Ruffles Original, Frito Lay, Casa Grande, AZ, USA), chocolate-covered peanut butter cups (Reese’s Minis Unwrapped, The Hershey Company, Hershey, PA, USA), and 11% weight/volume (w/v) high-fructose corn syrup-55 (HFCS) beverage (Best Flavors, Orange, CA, USA), placed in separate food/drink receptacles in the home cage for each animal. The 11% w/v concentration of HFCS was chosen to match the amount of sugar in sugar-sweetened beverages commonly consumed by humans, as we have previously modeled (*9, 35, 68*). CAF rats also received *ad libitum* access to water. Control (CTL) rats received the same number of food/drink receptacles, but they were filled with only standard chow (LabDiet 5001; PMI Nutrition International, Brentwood, MO, USA; 28.5% kcal from protein, 13.5% kcal from fat, 58.0% kcal from carbohydrate) or water, accordingly. Body weights and food intakes (including spillage collected on cardboard under the hanging wire cages) were measured three times per week between 8:30-10:30 (shortly before the onset of the dark cycle).

### General experimental overview

In order to match initial body weights by group in all cohorts, on PN 26 rats were pseudo-randomly assigned to groups receiving either the CAF diet or CTL diet from PN 26-56, which approximately represents the juvenile and adolescent periods in rats (*72, 73*). A generalized timeline of procedures is provided in Fig. 1A, and specific details for each experimental cohort can be found in Supplementary Methods and Materials.

### Metabolic assessments

All metabolic assessments were performed during the dark cycle between 11:00-14:00, and specific methos can be found in the Supplementary Materials and Methods file.

### Behavioral assessments

All behavioral assessments were performed during the dark cycle between 11:00-17:00.

#### Novel Location Recognition (NLR)

NLR was performed to assess spatial recognition memory, which relies on HPC function (*41, 42*). A grey opaque box (38.1 cm L × 56.5 cm W × 31.8 cm H), placed in a dimly lit room in which two adjacent desk lamps were pointed toward the floor, was used as the NLR apparatus. Rats were habituated to the empty apparatus and conditions for 10 min 1-2 days prior to testing. Testing constituted a 5-min familiarization phase during which rats were placed in the center of the apparatus (facing a neutral wall to avoid biasing them toward either object) with two identical objects placed in two corners of the apparatus and allowed to explore. The objects used were either two identical empty glass salt-shakers (that never contained salt) or two identical empty soap dispensers (that were never used with soap; first NLR time point), or two identical textured glass vases or two identical ceramic jugs (second NLR time point). Rats were then removed from the apparatus and placed in their home cage for 5 min. During this period, the apparatus and objects were cleaned with 10% ethanol solution and one of the objects was moved to a different corner location in the apparatus (i.e., the object was moved but not replaced). Rats were then placed in the center of the apparatus again and allowed to explore for 3 min. The types of objects used and the novel location placements were counterbalanced by group. The time each rat spent exploring the objects was quantified by hand-scoring of video recordings by an experimenter blinded to the animal group assignments and object exploration was defined as the rat sniffing or touching the object with the nose or forepaws.

#### Novel Object in Context (NOIC)

NOIC was conducted to assess HPC-dependent contextual episodic memory (*43, 44*). The 5-day NOIC procedure was adapted from previous research (*43*), and as conducted in previous work from our laboratory (*16, 35, 57, 74, 75*). Each day consisted of one 5-min session per animal, with cleaning of the apparatus and objects using 10% ethanol between each animal. Days 1 and 2 were habituation to the contexts – rats were placed in Context 1, a semitransparent box (41.9 cm L × 41.9 cm W × 38.1 cm H) with yellow stripes, or Context 2, a black opaque box (38.1 cm L × 63.5 cm W × 35.6 cm H). Each context was presented in a distinct room, both with similar dim ambient lighting yet with distinct extra-box contextual features. Rats were exposed to one context per day in a counterbalanced order per diet group for habituation. Following these two habituation days, on the next day each animal was placed in Context 1 containing single copies of Object A and Object B situated on diagonal equidistant markings with sufficient space for the rat to circle the objects (NOIC day 1). Objects were an assortment of hard plastic containers, tin canisters (with covers), and the Original Magic 8-Ball (two types of objects were used per experimental cohort time point; objects were distinct from what animals were exposed to in NOR and NLR). The sides where the objects were situated were counterbalanced per rat by diet group. On the following day (NOIC day 2), rats were placed in Context 2 with duplicate copies of Object A. The next day was the test day (NOIC day 3), during which rats were placed again in Context 2, but with single copies of Object A and Object B; Object B was therefore not a novel object, but its placement in Context 2 was novel to the rat. Each time the rats were situated in the contexts, care was taken so that they were consistently placed with their head facing away from both of the objects. On NOIC days 1 and 3, object exploration, defined as the rat sniffing or touching the object with the nose or forepaws, was quantified by hand-scoring of videos by an experimenter blinded to the animal group assignments. The discrimination index for Object B was calculated for NOIC days 1 and 3 as follows: time spent exploring Object B (the “novel object in context” in Context 2) / [time spent exploring Object A + time spent exploring Object B]. Data were then expressed as a percent shift from baseline as: [day 3 discrimination index – day 1 discrimination index] × 100. Rats with intact HPC function will preferentially explore the “novel object in context” on NOIC day 3, while HPC impairment will impede such preferential exploration (*43, 44*).

Methods for Novel Object Recognition (NOR), Zero Maze, and Open Field are found in Supplementary Methods and Materials.

#### Stereotaxic surgery

Stereotaxic surgery procedures are found in the Supplementary Methods and Materials file.

#### In vivo fiber photometry during NOIC

*Fiber photometry during NOIC procedure and data analyses.* Fiber photometry involves transmission of a fluorescence light through an optical patch cord (Doric Lenses, Quebec City, Quebec, Canada) and convergence onto the optic fibers implanted in the animals, which in turn emits neural fluorescence through the same optic fibers/patch cords and is focused on a photoreceiver. For the present experiments, animals were habituated to having patch cords attached to their HPC fiber optic cannulae prior to NOIC experimentation. Every animal had patch cords attached for both NOIC habituation days as well NOIC days 1-3, and unilateral photometry recordings were collected on NOIC days 1 and 3. Recording sessions lasted the duration of the trials on both NOIC days 1 and 3 (5 min each day per rat). Photometry signal was captured according to previous work by our group using the Neurophotometrics fiber photometry system (Neurophotometrics, San Diego, CA, USA) at a sampling frequency of 40 Hz with alternating wavelengths at 470 nm and 415 nm (*76, 77*). The signal representing ACh binding (470 nm) was corrected by subtracting the isosbestic signal (415 nm) and fitting the result to a biexponential curve using MATLAB (R202a, MathWorks, Inc., Natick, MA, USA). Corrected fluorescence signal was normalized within each recording session per rat by calculating ΔF/F using the average fluorescence signal for the entire recording. Resulting data was aligned with real-time location of each animal obtained from ANYmaze behavior tracking software (Stoelting, Wood Dale, IL, USA) using a customized MATLAB code. Investigations occurring at least 5 seconds apart were isolated for further analysis. Time-locked z-scores were obtained by normalizing ΔF/F for the 5 seconds following an object investigation episode to the ΔF/F at the onset of the investigation (time 0 s). Area under the curve was also calculated for z-scores for the 5 seconds following an object investigation episode. Aside from the patch cords and recording, NOIC was performed in the same manner as detailed above.

#### HPC ACh agonist infusion during NOIC

Animals were habituated to having their cannulae obturators removed and replaced one week prior to NOIC experimentation. Injectors for drug administration projected 1 mm beyond the guide cannulae for all infusions. NOIC procedures were carried out as described above with the exception that infusions of either ACh agonist or vehicle were performed in the animal housing room 3-8 minutes before each animal was placed in Context 2 on NOIC day 2 for the 5-min familiarization period with duplicate copies of Object A. This agonist administration timing was based off previous research (*78*) and pilot experiments in our lab that identified the consolidation period as critical to ACh function (data not shown). The broad, nonselective ACh agonist carbachol (0.8 μg total dose; 0.2 μL volume of 2 μg/μL per side; Y0000113, Millipore Sigma, St. Louis, MO, USA), the α7 nAChR specific agonist PNU 282987 (1.6 μg total dose; 0.2 μL volume of 4 μg/μL per side; PNU 282987, Tocris, Ellisville, MO, USA), or vehicle (artificial cerebrospinal fluid [aCSF] or 50/50% aCSF/dimethylsulfoxide [DMSO]) was infused bilaterally per rat. Volumes for bilateral HPC infusions were 200 nl/hemisphere (rate = 5 μL/min) via a 33-gauge injector and microsyringe attached to an infusion pump (Harvard Apparatus). Injectors were left in place for 20 s after infusions. Carbachol was dissolved in aCSF, and PNU 282987 was dissolved in 50/50% aCSF/DMSO. Following infusions, animals were carefully brought to the behavior room and proceeded with NOIC day 2 training as described above.

#### Tissue collection

Specific methodological details for tissue collection per experimental cohort can be found in the Supplementary Materials and Methods file.

#### Immunoblotting

Immunoblotting procedures can be found in Supplementary Materials and Methods.

#### Microbiome analyses

All microbiome analyses methods can be found in the Supplementary Materials and Methods file.

#### Statistical Analysis

Data are presented as means ± standard errors (SEM) for error bars in all figures. Each experimental group was analyzed with its respective control group per experimental cohort. Statistical analyses were performed using Prism software (GraphPad, Inc., version 8.4.2, San Diego, CA, USA) or R (4.2.0, R Core Team 2022. Significance was considered at *p* < 0.05. Detailed descriptions of the specific statistical tests per Fig. panel can be found in Table S1. In all cases, model assumptions were checked (normality by Shapiro-Wilk test and visual inspection of a qq plot for residuals per analysis, equal variance/homoscedasticity by an F test to compare variances and visual inspection of a residual plot per analysis). In one case (NLR time point 1 for cohort 1), data violated the assumption of homoscedasticity and thus a Welch’s correction was employed to account for the unequal variance. Group sample sizes were based on prior knowledge gained from extensive experience with rodent behavior testing and survival animal surgeries.

**List of Supplementary Materials**

Supplementary Methods and Materials

Fig. S1 to S7

Table S1

References 1-21

## One-sentence summary

This work reveals that consumption of a high-fat, high-sugar Western diet in early life leads to long-lasting memory impairments underscored by disruptions in hippocampus acetylcholine signaling.

## Acknowledgements

The authors would like to thank the Kanoski Lab undergraduate research assistants for their support with the behavioral and metabolic experiments.

## Author Contributions

Conceived and designed the study: AMRH, LT, SEK

Maintained subjects on the dietary model: AMRH, AEK, MEK, LT, CG, NT, AA

Performed metabolic assessments: AMRH, AEK, LT

Performed behavioral experiments: AMRH, LTL, AEK, LT, NT, AA

Performed stereotaxic surgeries: AMRH, JJR, KSS, LDS

Established in vivo photometry procedures and performed photometry data analyses: LTL, KSS, LDS, AMRH

Carried out tissue collection: AMRH, AEK, MEK, LT, JJR Performed immunoblot experiments: KND

Performed microbiome analyses and machine learning analyses: SS, AAF

Performed all other statistical analyses: AMRH, SEK

Writing – original draft: AMRH, SEK

Writing – review & editing: LTL, AEK, SS, MEK, LT, JJR, KSS, CG, NT, AA, KND, LDS, AAF

## Data and materials availability

All data associated with this work are present in the paper or the Supplementary Materials. The 16s rRNA microbiome sequencing data will be made available in the National Center for Biotechnology Information (NCBI) Sequence Read Archive (SRA).

## Supplementary Materials for

### Dietary model – part 2

The percentage of total kilocalories consumed from each of the CAF diet components was calculated by multiplying the measured weights of food/drink consumed per rat by the energy density of each component (4.7 kcal/g for HFHS diet, 5.7 kcal/g for potato chips, 5.1 kcal/g for peanut butter cups, 0.296 kcal/g for HFCS beverage), and in the same manner the total energy intake for the CTL group was calculated using the energy density of standard chow (3.36 kcal/g). The percentage of kilocalories consumed from each macronutrient (% kcal from carbohydrate, % kcal from protein, % kcal from fat) for the CAF diet groups was calculated using the kilocalories consumed from each CAF diet component and each component’s macronutrient composition.

Experimental cohort 1: Early life cafeteria (CAF) diet followed by healthy diet intervention: behavioral and metabolic outcomes. Rats (n=24 total) received either the CAF or CTL diet from PN 26-86 (n=12 per group). The 30-day early life diet period encompassed PN 26-56, after which point behavioral testing occurred while rats remained on their respective diets (Fig. 1A). Behavioral tests began with Novel Object Recognition (NOR) and Novel Location Recognition (NLR) experiments conducted in a counterbalanced fashion by diet group on PN 60-61 (with habituation on PN 59). Rats then were tested in the Novel Object in Context (NOIC) task from PN 68-70 (with habituation on PN 66 and 67). Zero Maze and Open Field tests were performed on PN 74 and PN 75, respectively. An intraperitoneal glucose tolerance test (IPGTT) was conducted on PN 77, and rats on the CAF diet were switched to the standard healthy chow diet (i.e., the same diet that the CTL group received throughout the study) on PN 86. All the outcomes measured up to PN 86 are considered the assessments “before the healthy diet intervention.” The 30-day healthy diet intervention period for the CAF rats lasted from PN 86-116. For the outcomes assessed after the healthy diet intervention, NOIC was performed from PN 125-127 (habituation on PN 123 and 124), body composition was analyzed on PN 132, NOR was performed on PN 136, Zero Maze was conducted on PN 140, NLR was assessed on PN 143, and an IPGTT was performed on PN 147. Fecal samples were collected for 16s rRNA sequencing analyses of the gut microbiome on PN 56 and PN 116. Tissue was harvested on PN 150.

Experimental cohort 2: Early life CAF diet without healthy diet intervention: tissue collection. Rats in this cohort (n=24 total) received either CAF diet or standard chow diet as control (CTL) from PN 26-74 (n=12 per group), with NOIC testing from PN 68-70 (with habituation on PN 66 and 67) followed by tissue harvest at PN 74. Through this cohort, we were able to collect cecal content (which is a terminal endpoint) following CAF exposure without a healthy diet intervention.

Experimental cohort 3: Early life CAF diet followed by healthy diet intervention: in vivo fiber photometry ACh binding analyses. Rats (n=24 total) received either CAF diet or standard chow diet as control (CTL) from PN 26-56 (n=12 per group). The 30-day early life diet period encompassed PN 26-56, as in previous cohorts, but the CAF group was switched onto the healthy standard chow at PN 56 to mark the beginning of the healthy diet intervention period. Body composition measures were also performed on PN 56.

Stereotaxic surgeries for in vivo fiber photometry analyses were performed on PN 67, PN 71, and PN 72 (n=8 rats per day, counterbalanced per group). This timing was selected to allow the CAF rats to acclimate to the standard healthy chow for 10+ days before surgery as well as for adequate recovery time before behavior testing while also accommodating for 30+ days of the healthy diet intervention period before behavior testing. NOIC was performed in squads from PN 95-125. Rats from this cohort were transcardially perfused on PN 226 and brain tissue was harvested for histological confirmation of AAV injection placement.

Experimental cohort 4: Early life CAF diet followed by healthy diet intervention: ACh pharmacology. Rats in this cohort (n=30 total) all received CAF diet from PN 26-56, but they were assigned to different groups at PN 26 according to which ACh agonist they would receive during NOIC behavior testing (n=10 per group). The 30-day early life diet period encompassed PN 26-56, as in previous cohorts, but the CAF group was switched onto the healthy standard chow at PN 56 to mark the beginning of the healthy diet intervention period. Stereotaxic surgeries to install bilateral indwelling cannulae into the dorsal HPC were performed on PN 66-69 (n=8 rats per day, counterbalanced per group). This surgery timing was selected to allow the CAF rats to acclimate to the standard healthy chow for 10+ days before surgery as well as for adequate recovery time before behavior testing while also accommodating for 30+ days of the healthy diet intervention period before neuropharmacological behavior testing. NOIC was performed from PN 94-96 (habituation on PN 92 and 93). Rats from this cohort were infused with blue sky ink through their cannulae on PN 109 as described below and immediately euthanized to enable validation of infusion sites.

### Metabolic assessments

#### Body composition

Nuclear magnetic resonance (NMR) was used to quantify body composition as previously described (1, 2) both immediately after the 30-d CAF diet period in early life (PN 56) as well as after the 30-d healthy diet intervention in adulthood (PN 132). Briefly, rats were food-restricted for 1 h, weighed, and scanned using the Bruker NMR Minispec LF 90II (Bruker Daltonics, Inc., Billerica, MA, USA) interfaced to a computer equipped with Minispec Plus 4.1.5 software. This system employs time-domain NMR signals from all protons to quantify body composition and is advantageous because it is non-invasive and does not require administration of anesthesia. The ratio of fat-to-lean mass per subject was calculated as [fat mass (g) / lean mass (g)].

#### Glucose tolerance

An intraperitoneal glucose tolerance test (IPGTT) was performed to examine glucose metabolism both immediately after the 30-d CAF diet period in early life (PN 77) as well as after the 30-d healthy diet intervention in adulthood (PN 147). Rats were food-restricted 22 h prior to the test, following established procedures (2–4). Blood was sampled from the tip of the tail for baseline blood glucose readings (fasted state) and all subsequent blood glucose readings and measured using a glucometer (One Touch Ultra2, LifeScan, Inc., Milpitas, CA, USA). Following the baseline reading, each rat received an intraperitoneal (IP) injection of a 50% glucose solution (1 g/kg body weight; Dextrose anhydrous, VWR Chemicals, LLC, Solon, OH, USA). Aside from the baseline reading (collected immediately before injection), blood glucose readings were subsequently obtained at the 30, 60, 90, and 120 min time points following injection. Area under the curve was calculated using the trapezoidal rule with 0 mg/dL set as the mathematical “baseline”.

### Behavioral assessments – part 2

#### Novel Object Recognition (NOR)

NOR was used to evaluate exploration of novelty in the form of a novel object, which under testing parameters employed is perirhinal cortex-dependent and not HPC-dependent (5). A grey opaque box (38.1 cm L × 56.5 cm W × 31.8 cm H), placed in a dimly lit room in which two adjacent desk lamps were pointed toward the floor, was used as the NOR apparatus. Procedures followed as described in (6), modified from (7). Rats were habituated to the empty apparatus and conditions for 10 min 1-2 days prior to testing. The test consisted of a 5-min familiarization phase during which rats were placed in the center of the apparatus (facing a neutral wall to avoid biasing them toward either object) with two identical objects and allowed to explore. The objects used were either two identical cans or two identical stem-less wine glasses (first NOR time point), or two identical tin canisters (with tin covers) or two identical ceramic vases (second NOR time point). Rats were then removed from the apparatus and placed in their home cage for 5 min. During this period, the apparatus and objects were cleaned with 10% ethanol solution and one of the objects was replaced with a different one (for time point 1: either the can or glass – whichever the animal had not previously been exposed to – i.e., the “novel object”; for time point 2: either the tin canister or vase – whichever the animal had not previously been exposed to – i.e., the “novel object”). Rats were then placed in the center of the apparatus again and allowed to explore for 3 min. The novel object and side on which the novel object was placed were counterbalanced per treatment group. The time each rat spent exploring the objects was quantified by hand-scoring of video recordings by an experimenter blinded to the animal group assignments. Object exploration was defined as the rat sniffing or touching the object with the nose or forepaws.

#### Zero Maze

Following established procedures (2, 3, 6), the Zero Maze procedure was used to examine anxiety-like behavior. The Zero Maze apparatus consisted of an elevated circular track (11.4 cm wide track, 73.7 cm height from track to the ground, 92.7 cm exterior diameter) that is divided into four equal length segments: two sections with 3-cm high curbs (open), and two sections with 17.5 cm high walls (closed). Ambient lighting was used during testing. Rats were placed in the maze on an open section of the track and allowed to roam for 5 min, during which time they could freely ambulate through the different segments of the track. The apparatus was cleaned with 10% ethanol between rats. The time each rat spent in the open segments of the track (defined as the center of the rat in an open arm) as well as the number of entries into the open segments of the track were measured via video recording using ANYmaze activity tracking software (Stoelting Co., Wood Dale, IL, USA).

#### Open Field

Open Field was performed to test for locomotor activity as well as anxiety-like behavior following previous procedures (4, 6, 8, 9). The Open Field apparatus was a gray arena (53.5 cm L × 54.6 cm W × 36.8 H) with a designated center zone within the arena (19 cm × 17.5 cm). The center zone was maintained under diffused lighting (44 lux) compared to the corners and edges (∼30 lux). For testing, rats were placed in the center of the apparatus and allowed to freely explore for 10 min. The apparatus was cleaned with 10% ethanol between rats. Video recording using ANYmaze activity tracking software (Stoelting Co., Wood Dale, IL, USA) was used to quantify time spent in the center zone (a measuring of anxiety-like behavior) and distance travelled in the apparatus during the task (locomotor activity).

### Stereotaxic surgery

#### Intracranial viral injection and in vivo photometry optic fiber placement

Rats were anesthetized and sedated via intramuscular injections of a cocktail of ketamine (90.1 mg/kg body weight), xylazine (2.8 mg/kg body weight), and acepromazine (0.72 mg/kg body weight), followed by a pre-operative subcutaneous injection of analgesic (buprenorphine SR, 1.0 mg/kg body weight). Upon sedation, the surgical site was shaved and prepared with iodine and ethanol swabbing. The animals were then placed in a stereotaxic apparatus. The iAChSnFR adeno-associated virus (AAV; 1 μL; original titer ≥ 7×10¹² vg/mL diluted 1:2 with artificial cerebral spinal fluid; pAAV.hSynap.iAChSnFR, 137950-AAV1, Addgene, Watertown, MA, USA), validated as a selective in vivo ACh sensor in rodents (10–14), was delivered into the dorsal dentate gyrus of the HPC (anterior/posterior [A/P]: -3.24; medial/lateral [M/L]: +/-1.80; dorsal/ventral [D/L]: -3.50; with 0 as the reference point at bregma for A/P, M/L, and D/L) using a microinfusion pump (Harvard Apparatus, Cambridge, MA, USA) equipped with a 33-gauge microsyringe injector connected to a PE20 catheter and Hamilton syringe. The injection flow rate was set to 5 μL/min. To allow for complete delivery of infusate, injectors were kept in place for 2 min post-injection. Fiber optic cannulae (flat 400 μm core, 0.48 numerical aperture, 5mm; Doric Lenses Inc., Quebec, Canada) were implanted in the dorsal dentate gyrus at the same coordinates as the viral infusion (A/P: -3.24; M/L: +/-1.80; D/L: -3.50; with 0 as the reference point at bregma for A/P, M/L, and D/L). Optic fibers were then affixed to the skull with jeweler’s screws, instant adhesive super glue, and dental cement.

#### Bilateral indwelling cannulae in HPC

Intramuscular injections of a cocktail of ketamine (90.1 mg/kg body weight), xylazine (2.8 mg/kg body weight), and acepromazine (0.72 mg/kg body weight), followed by a pre-operative subcutaneous injection of analgesic (buprenorphine SR, 1.0 mg/kg body weight) were administered to anesthetize and sedate rats. The surgical site was shaved and prepared with iodine and ethanol swabbing, and subsequently the animals were situated in a stereotaxic apparatus. Indwelling bilateral guide cannulae (26-gauge, Plastics One, Protech International Inc., Boerne, TX, USA) were then placed targeting the dorsal dentate gyrus of the HPC (A/P: -4.08; M/L: +/-2.50; D/L: -2.60; with 0 as the reference point at bregma for A/P, M/L, and D/L). Cannulae were fastened to the skull with jeweler’s screws, instant adhesive super glue, and dental cement. Obturators were secured in the cannulae to prevent blockage until experimentation.

### Tissue collection

#### Experimental cohorts 1 and 2

Following intramuscular injection of a cocktail of ketamine (90.1 mg/kg body weight), xylazine (2.8 mg/kg body weight), and acepromazine (0.72 mg/kg body weight), animals were rapidly decapitated. Cecal content was collected as described below, and brains were flash-frozen in a beaker containing -30°C isopentane, surrounded by dry ice. All samples were stored at -80°C until further processing and analysis. Tissue punches of the dorsal HPC (atlas levels 28-32; one 2 mm diameter punch in the CA1 / dentate gyrus, one 1 mm diameter punch in the CA3) for immunoblot analyses were collected using the Leica CM cryostat (Wetzlar, Germany) and anatomical landmarks were based on the Swanson rat brain atlas (15). Punches were stored at -80°C until used for later immunoblotting assays as described below.

#### Experimental cohort 3

Rats were anesthetized via an intramuscular injection of a ketamine (90.1 mg/kg body weight), xylazine (2.8 mg/kg body weight), and acepromazine (0.72 mg/kg body weight) cocktail and then transcardially perfused with 0.9% sterile saline (pH 7.4) followed by 4% paraformaldehyde (PFA) in 0.1 M borate buffer (pH 9.5). Brains were dissected from the skull and post-fixed in PFA with 15% sucrose for 24 h, and then flash-frozen in isopentane cooled in dry ice. Brains were then stored at -80°C. On a freezing microtome, brains were sectioned to 30 μm thickness and sections were collected in 5-series, stored in antifreeze solution at -20°C, mounted, and coverslipped using 50% glycerol in 0.02 M KPBS, with edges sealed using clear nail polish. Photomicrographs for confirmation of AAV injection sites were captured using a Nikon 80i camera (Nikon DSQI1, 1280X1024 resolution, 1.45 megapixel).

#### Experimental cohort 4

Injection of a cocktail of ketamine (90.1 mg/kg body weight), xylazine (2.8 mg/kg body weight), and acepromazine (0.72 mg/kg body weight) was administered intramuscularly. Pontamine sky blue ink (2%, 200 nL per hemisphere) was then infused into the indwelling HPC bilateral cannulae to allow for postmortem verification of the infusion sites. Thereafter, rats were rapidly decapitated. Brains were extracted and post-fixed in 10% formalin until further processing. Cannulae placements were examined through anatomical verification of the position of the pontamine sky blue ink infusions. Animals with ink confined to the dorsal HPC based on the Swanson rat brain atlas (15) in one or both cannulae were included in data analysis. No animals were excluded from data analysis based on this criterion.

### Immunoblotting (experimental cohort 1)

Dorsal HPC tissue punches from experimental cohort 1 were analyzed for levels of ChAT, VAChT, and AChE. Proteins in brain lysates were separated using sodium dodecyl sulfate polyacrylamide gel electrophoresis, transferred onto poly-vinylidene difluoride membranes, and subjected to enhanced chemiluminescence for immunodetection analysis (Chemidoc XRS, BioRad). A rabbit polyclonal anti-choline acetyltransferase antibody (1:100; Sigma-Aldrich, catalog # AB143) was used to determine the concentration of ChAT relative to a load control signal detected by a rabbit anti-β-tubulin antibody (1:1000; Cell Signaling Technology, 9F3 rabbit mAb, catalog # 2128). A rabbit polyclonal anti-vesicular ACh antibody (1:1000; Sigma-Aldrich, catalog # ABN100) was used to determine the concentration of VAChT relative to a load control signal detected by a rabbit anti-β-actin antibody (1:1000; Abcam, catalog # ab8227). A rabbit recombinant anti-acetylcholinesterase antibody (1:1000; Abcam, catalog # ab183591) was used to determine the concentration of AChE relative to a loading control signal detected by a rabbit anti-β-actin antibody (1:1000; Abcam, catalog # ab8227). Blots were quantified with densitometry analysis using ImageJ as previously reported (8, 16, 17).

### Microbiome analyses

#### Fecal collection

Fecal collections were performed on PN 56 and 116 from the first experimental cohort. Rats were placed into a sterile cage (without bedding) and mildly restrained until defecation. Two fecal samples per rat per time point were collected. Each was weighed immediately upon defecation under sterile conditions and placed into a distinct Dnase/Rnase-free 2 mL cryogenic vial and immediately embedded in dry ice (one sample per vial). All samples were then stored in a -80°C freezer until further analysis. Between every rat, all cages and sample collection materials were thoroughly cleaned with 70% ethanol. All fecal collections were performed between 12:00 and 15:00 (during the dark cycle).

#### Cecal content collection

Collecting cecal content is a terminal procedure, and thus it was performed on the terminal tissue collection days for the first two experimental cohorts (PN 150 and PN 74, respectively). Following euthanasia procedures (described above), an abdominal incision was made to expose the intestines. The cecum was identified and a small incision was made therein. Cecal content was then collected directly into a pre-tared Dnase/Rnase-free 2 mL cryogenic vial, weighed, and immediately embedded into dry ice. All samples were then stored in a -80°C freezer until further analysis. Two vials of cecal content were collected per rat. Between every rat, all cecal content collection materials were thoroughly cleaned with 70% ethanol. Cecal content collections were performed between 11:00 and 17:00 (during the dark cycle).

#### Sample processing and 16S rRNA sequencing

Bacterial genomic DNA was extracted from fecal samples using Laragen Inc.’s validated in-house fecal DNA extraction protocol. Quantification of 16S rRNA gene loads was performed by qPCR using the SYBR Green master mix (Roche, Basel, Switzerland), universal 16S rRNA gene primers55, and the QuantStudio 5 thermocycler (cycling parameters: 2 min at 50 °C, 10 min at 95 °C, 40 cycles of 10 s at 95 °C, and 45 s at 62 °C). Sequencing libraries were generated using previously established methods (18). The V4 regions of the 16S rRNA gene were PCR amplified using individually barcoded universal primers and 3 ng of the extracted genomic DNA. The PCR reaction was set up in a single reaction, and the PCR product was purified using Laragen Inc.’s validated in-house bead-based purification. Purified PCR product from each sample (250 ng) was pooled and sequenced by Laragen, Inc., using the Illumina MiSeq platform and 2 × 150 bp reagent kit for paired-end sequencing.

#### Microbiome data processing and analysis

The 16S rRNA amplicon sequences were analyzed with DADA2 and QIIME2 following the developers’ instructions (19, 20). The forward reads were truncated to 120 bp and denoised with DADA2 to 2,162 amplicon sequence variants (ASV). Chimeras were removed with the ‘consensus’ method using DADA2. The ASVs were classified with ‘classify-sklearn’ in QIIME2 using the SILVA database (release 138). The taxonomic abundance tables were normalized as previously described (21) to correct for different sequencing depth across samples. Statistical analyses were performed with R (4.2.0). The Bray-Curtis dissimilarity between samples were calculated using the genus level abundance and visualized using the principal coordinates analysis (PCoA) with function ‘capscale’ in R package ‘vegan’. The associations of microbial community profiles and diet were analyzed with the PERMANOVA test using the function ‘adonis’ in the same package. Shannon index was used for analyzing alpha-diversity.

Wilcoxon rank sum test was used to analyze the associations of individual taxa and diet. Taxa with presence in less than 25% samples were excluded to avoid spurious associations. P-values were adjusted with the Benjamini-Hochberg method for multiple hypotheses testing. The significant taxa (FDR<0.1) were highlighted in the phylogenetic tree using R package ‘plotmicrobiome’. The associations of taxa and NOIC were analyzed with Spearman’s correlations.

#### Machine learning – predicting NOIC performance using microbiome

Machine learning approaches were used to test if NOIC memory performance measures could be predicted from gut microbiome profiles after the WD access period but before the healthy diet intervention. Random forest regression models were built with R package ‘randomForest’ to predict NOIC and NLR from the taxonomic abundance at the genus level. Performance of the models were evaluated with 3-fold and 4-fold cross validation.

---

**Supplementary Figure S1.**
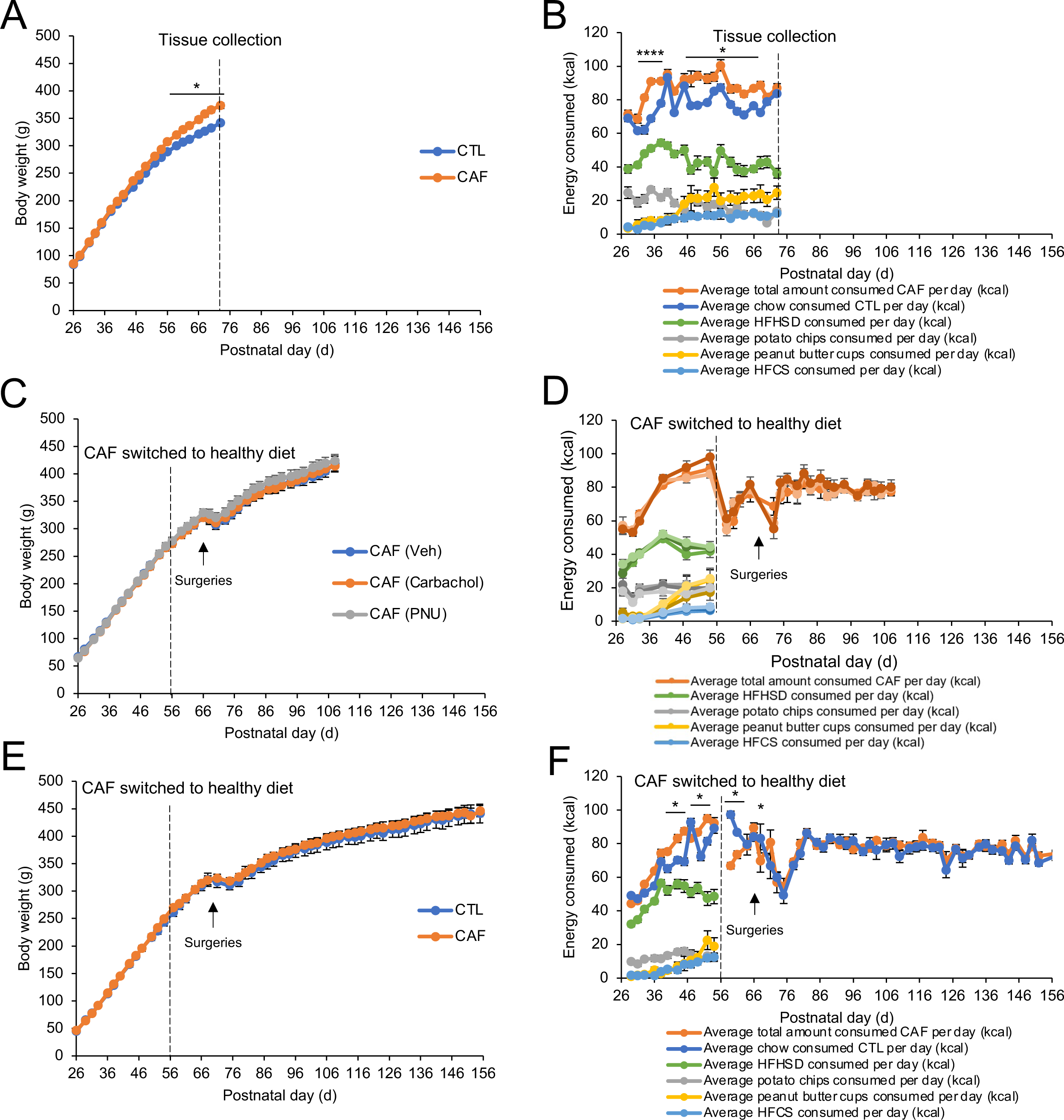
Body weight and energy intake of subsequent experimental cohorts. (A) Body weight over time for tissue cohort with no healthy diet intervention. (B) Energy consumption over time for tissue cohort with no healthy diet intervention. (C) Body weight over time for acetylcholine receptor agonist cohort. (D) Energy consumption over time for acetylcholine receptor agonist cohort. (E) Body weight over time for in vivo fiber photometry cohort. (F) Energy consumption over time for in vivo fiber photometry cohort. CAF, cafeteria diet group; CTL, control group; HFCS, high-fructose corn syrup; HFHSD, high-fat high-sugar diet. Error bars represent ± SEM. *P<0.05, ****P<0.0001.

**Supplementary Figure S2.**
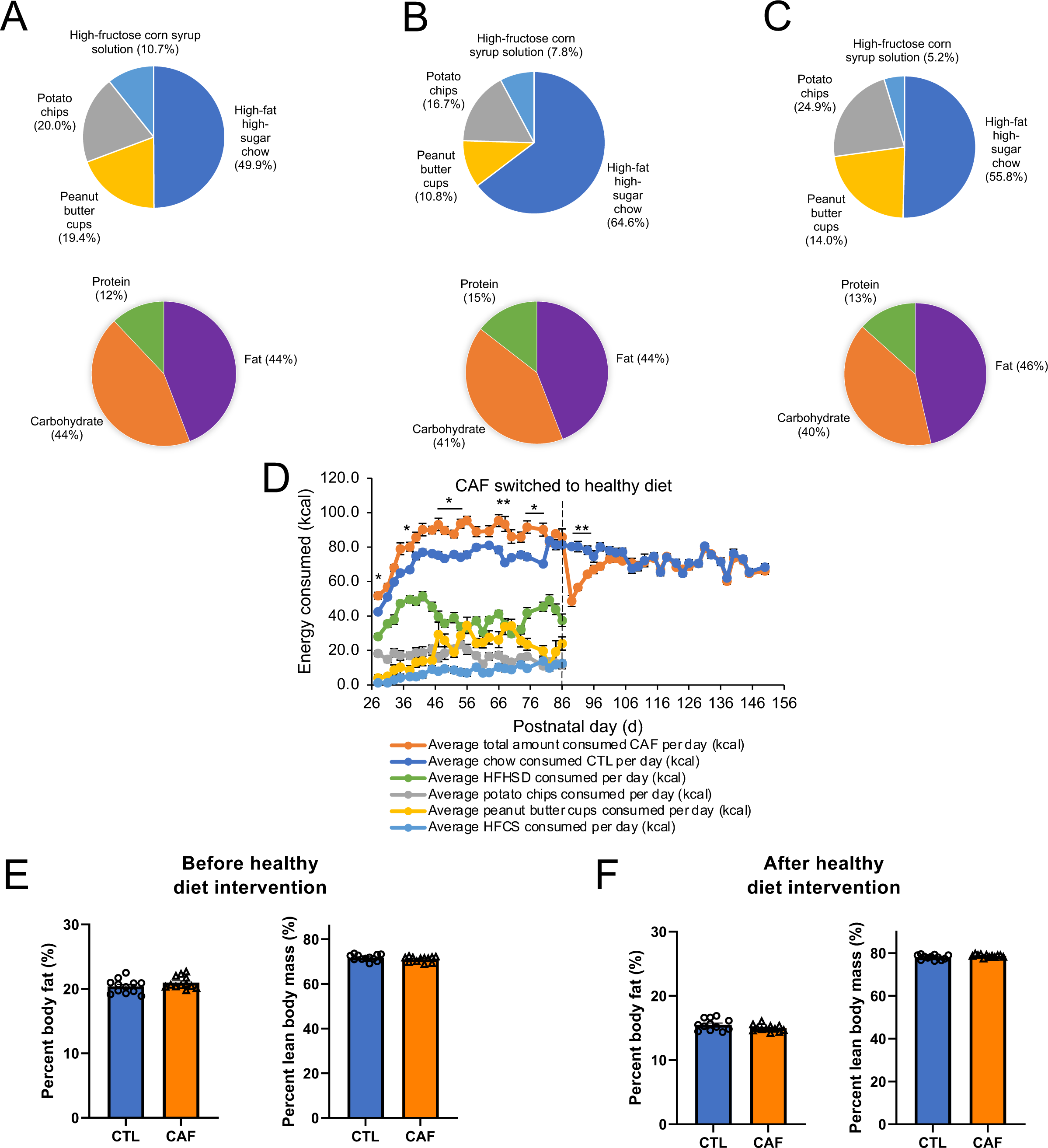
(A-C) Breakdown of percentages of kilocalories consumed from the cafeteria diet components and macronutrients reveals consistency across cohorts. (E-F) No differences were observed for percent body fat or lean mass between CTL and CAF groups both before and after the healthy diet intervention period. CAF, cafeteria diet group; CTL, control group; HFCS, high-fructose corn syrup; HFHSD, high-fat high-sugar diet. Error bars represent ± SEM.

**Supplementary Figure S3.**
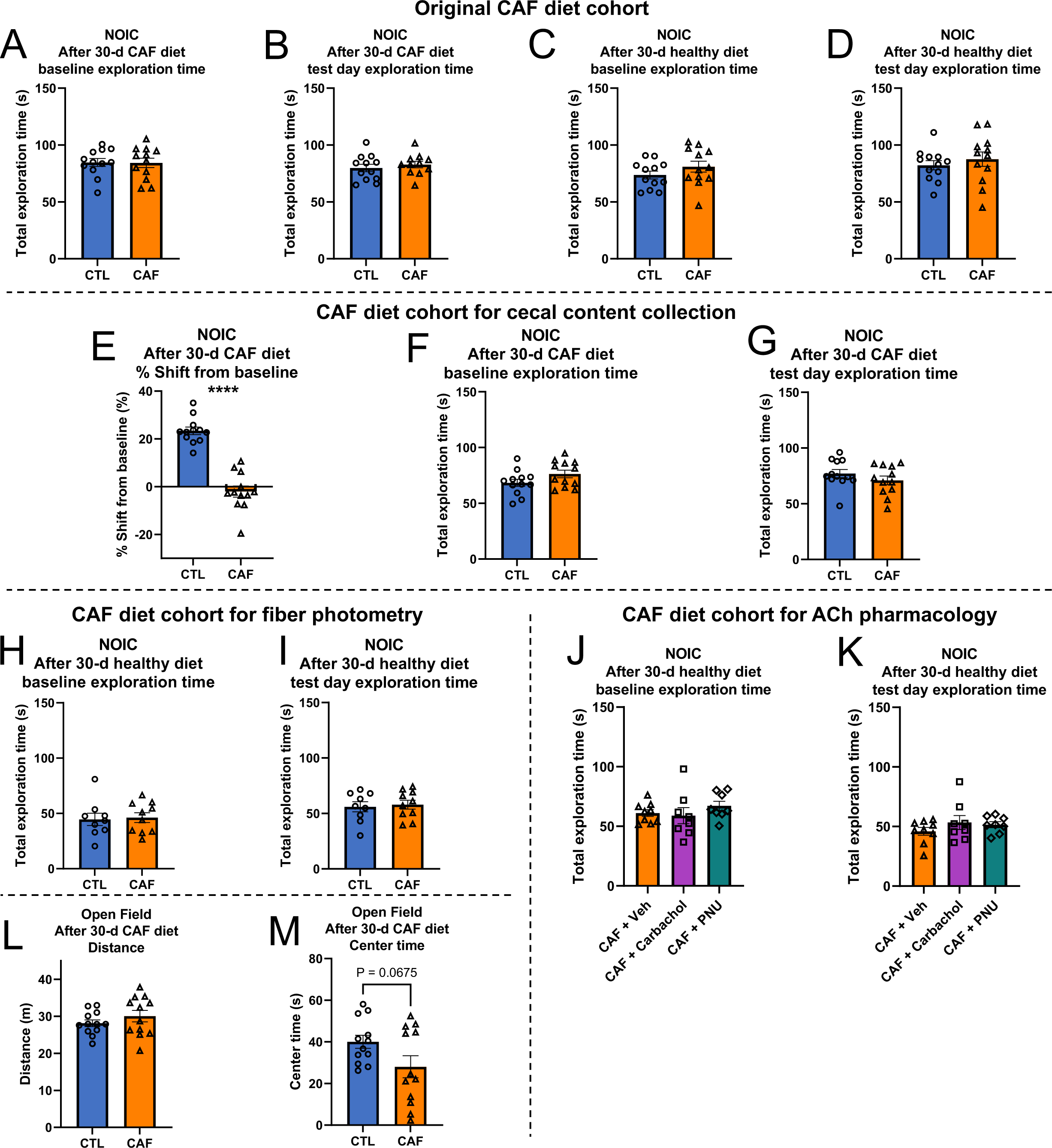
(A-D, F-K) No differences in total object exploration time between the early life Western diet and control groups across cohorts, and (E) replication of hippocampal-dependent memory impairment in CAF cohort for tissue collection. (L-M) No differences observed in anxiety-like behavior and locomotor activity. CAF, cafeteria diet group; CTL, control group; Carbachol, general ACh agonist; NOIC, novel object in context; PNU, α7 nicotinic receptor agonist. Error bars represent ± SEM. ****P<0.0001.

**Supplementary Figure S4.**
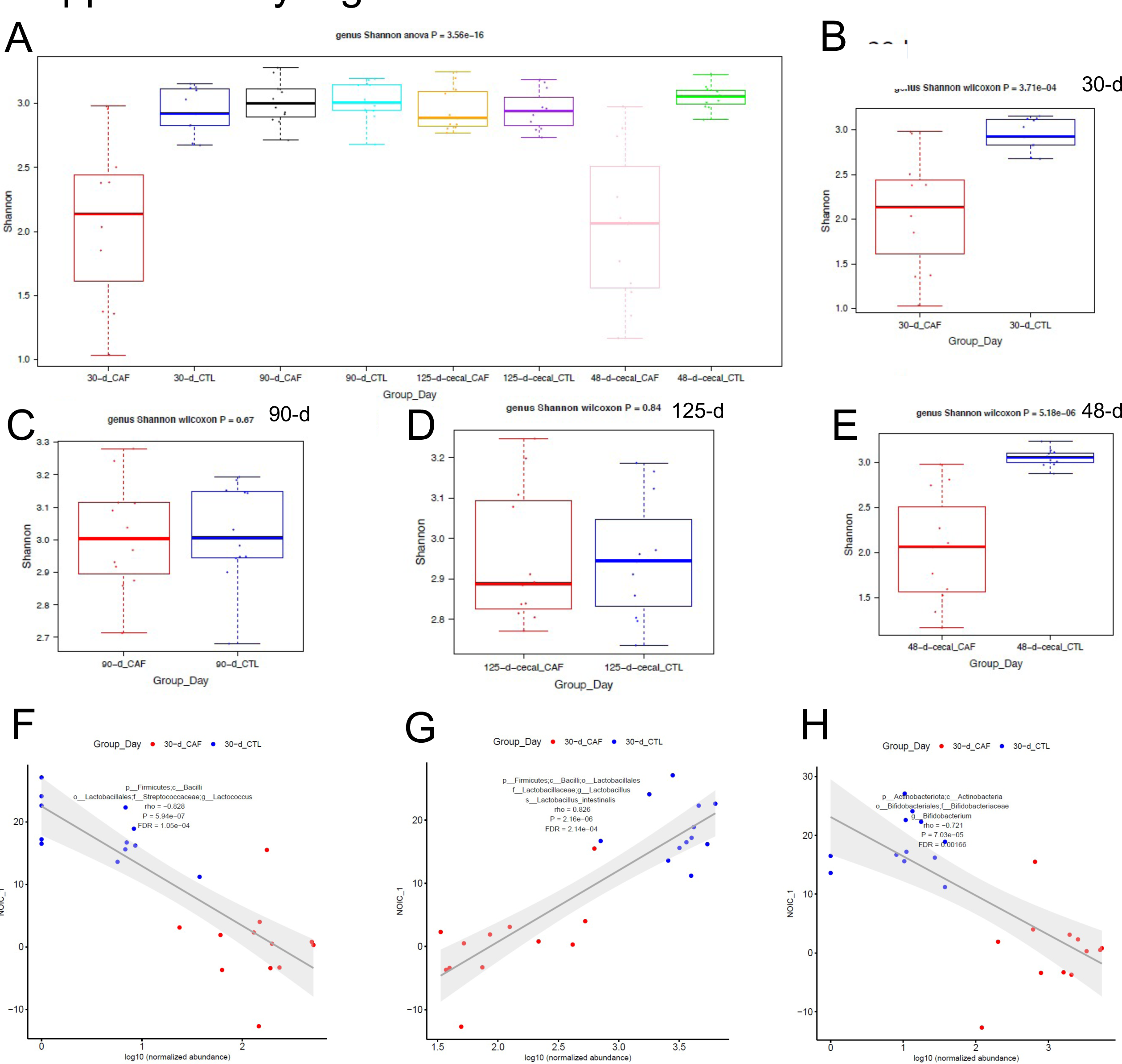
Microbial diversity measures across time. (A) Shannon diversity indices in CAF vs. CTL groups at different time point. (B-E) Shannon diversity indices in CAF and CTL groups split per time point. (F-H) Correlational analyses between specific bacterial taxa and contextual episodic memory performance after the 30-d CAF diet period (but prior to the healthy diet intervention). CAF, cafeteria diet group; CTL control diet group; NOIC, Novel Object in Context. P-values are indicated in each subpanel.

**Supplementary Figure S5.**
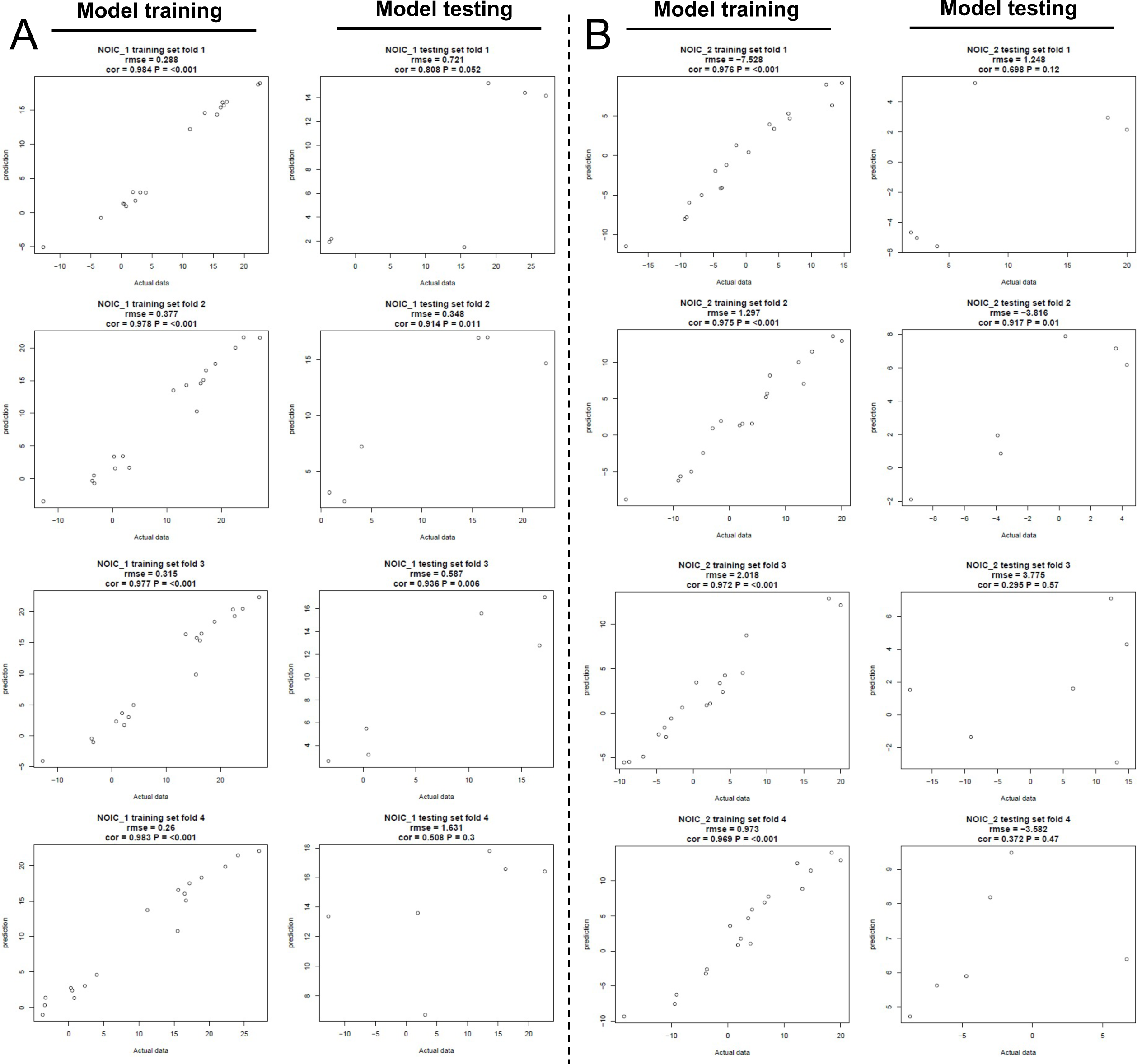
Machine learning analyses further reveal that microbial taxonomic composition following the Western diet period does not predict memory performance after the heathy diet intervention period. (A) Machine learning analysis testing prediction of memory performance in NOIC before the healthy diet intervention period. (B) Machine learning analysis testing prediction of memory performance in NOIC after the healthy diet intervention period. CAF, cafeteria diet group; CTL control diet group; NOIC, novel object in context. P-values are indicated in each sub-panel.

**Supplementary Figure S6.**
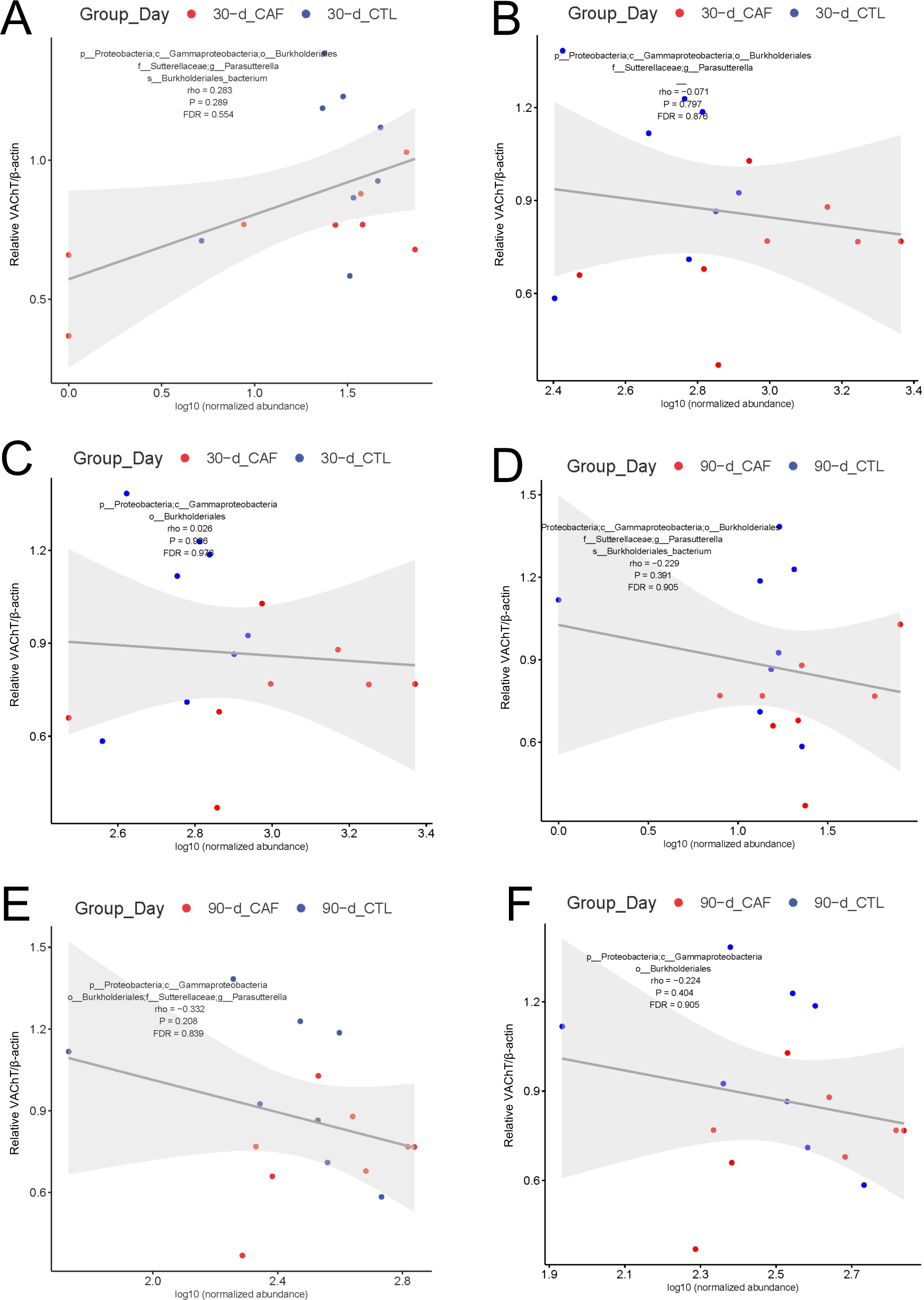
(A-F) Correlations between microbial taxa and VAChT levels from immunoblotting before the healthy diet intervention period (30-d; A-C) and after the healthy diet intervention period (90-d; D-F). CAF, cafeteria diet group; CTL control diet group. P-values are indicated in each sub-panel.

**Supplementary Figure S7.**
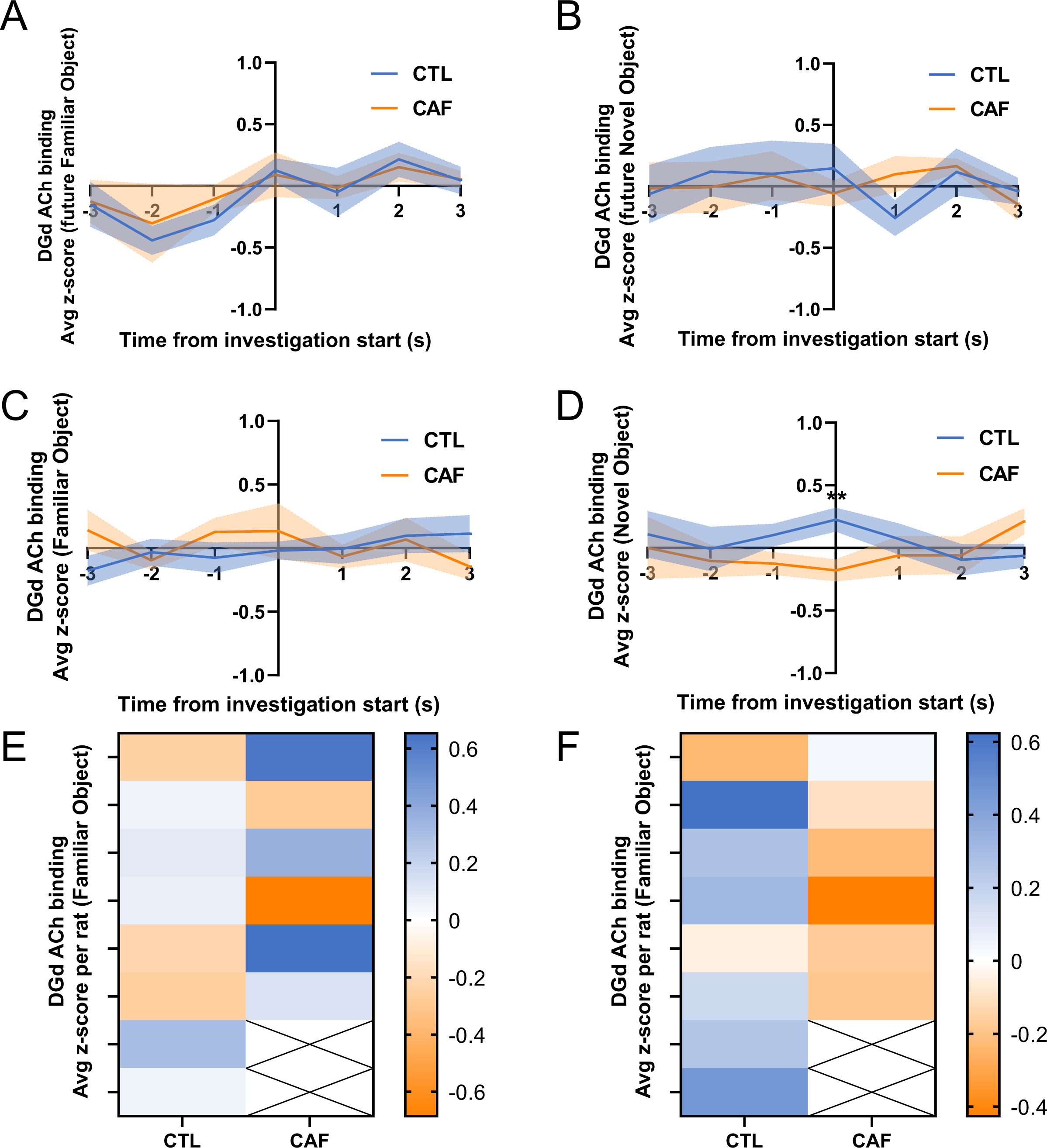
Photometry analyses for hippocampus acetylcholine (ACh) binding time-locked to investigations for baseline and test day object exploration in the contextual episodic memory task [Novel Object in Context, NOIC]. (A) ACh binding over time during day 1 of NOIC for the future familiar object in context. (B) ACh binding over time during day 1 of NOIC for the future novel object in context. (C) ACh binding over time during day 3 of NOIC for the familiar object in context. (D) ACh binding over time during day 3 of NOIC for the novel object in context. (E) Heat map of average z-score per rat at the initiation of an investigation for the familiar object in context (0 s). (F) Heat map of average z-score per rat at the initiation of an investigation for the novel object in context (0 s). ACh, acetylcholine; CAF, cafeteria diet group; CTL control diet group; NOIC, Novel Object in Context. Shading represents ± SEM for average values over time. **P<0.01.

## References

1. Christopher R. Wilson, Mai K. Tran, Katrina L. Salazar, Martin E. Young, H. Taegtmeyer, Western diet, but not high fat diet, causes derangements of fatty acid metabolism and contractile dysfunction in the heart of Wistar rats. Biochemical Journal 406, 457–467 (2007).

2. C. A. Shively, S. E. Appt, M. Z. Vitolins, B. Uberseder, K. T. Michalson, M. G. Silverstein-Metzler, T. C. Register, Mediterranean versus Western Diet Effects on Caloric Intake, Obesity, Metabolism, and Hepatosteatosis in Nonhuman Primates. Obesity 27, 777–784 (2019).

3. I. Heinonen, P. Rinne, S. T. Ruohonen, S. Ruohonen, M. Ahotupa, E. Savontaus, The effects of equal caloric high fat and western diet on metabolic syndrome, oxidative stress and vascular endothelial function in mice. Acta Physiologica 211, 515–527 (2014).

4. Y. Luo, C. M. Burrington, E. C. Graff, J. Zhang, R. L. Judd, P. Suksaranjit, Q. Kaewpoowat, S. K. Davenport, A. M. O’Neill, M. W. Greene, Metabolic phenotype and adipose and liver features in a high-fat Western diet-induced mouse model of obesity-linked NAFLD. American Journal of Physiology-Endocrinology and Metabolism 310, E418–E439 (2016).

5. S. E. Kanoski, T. L. Davidson, Western diet consumption and cognitive impairment: Links to hippocampal dysfunction and obesity. Physiology & Behavior 103, 59–68 (2011).

6. H. Francis, R. Stevenson, The longer-term impacts of Western diet on human cognition and the brain. Appetite 63, 119–128 (2013).

7. E. E. Noble, T. M. Hsu, S. E. Kanoski, Gut to Brain Dysbiosis: Mechanisms Linking Western Diet Consumption, the Microbiome, and Cognitive Impairment. Frontiers in Behavioral Neuroscience 11, (2017).

8. L. Tsan, L. Décarie-Spain, E. E. Noble, S. E. Kanoski, Western Diet Consumption During Development: Setting the Stage for Neurocognitive Dysfunction. Frontiers in Neuroscience 15, (2021).

9. T. M. Hsu, V. R. Konanur, L. Taing, R. Usui, B. D. Kayser, M. I. Goran, S. E. Kanoski, Effects of sucrose and high fructose corn syrup consumption on spatial memory function and hippocampal neuroinflammation in adolescent rats. Hippocampus 25, 227–239 (2015).

10. M. D. Kendig, R. A. Boakes, K. B. Rooney, L. H. Corbit, Chronic restricted access to 10% sucrose solution in adolescent and young adult rats impairs spatial memory and alters sensitivity to outcome devaluation. Physiology & Behavior 120, 164–172 (2013).

11. T. L. Davidson, A. Monnot, A. U. Neal, A. A. Martin, J. J. Horton, W. Zheng, The effects of a high-energy diet on hippocampal-dependent discrimination performance and blood–brain barrier integrity differ for diet-induced obese and diet-resistant rats. Physiology & behavior 107, 26–33 (2012).

12. S. E. Kanoski, Y. Zhang, W. Zheng, T. L. Davidson, The effects of a high-energy diet on hippocampal function and blood-brain barrier integrity in the rat. Journal of Alzheimer’s Disease 21, 207–219 (2010).

13. L. E. Jarrard, On the role of the hippocampus in learning and memory in the rat. Behavioral and Neural Biology 60, 9–26 (1993).

14. C. M. Bird, N. Burgess, The hippocampus and memory: insights from spatial processing. Nature Reviews Neuroscience 9, 182–194 (2008).

15. K. Henke, B. Weber, S. Kneifel, H. G. Wieser, A. Buck, Human hippocampus associates information in memory. Proceedings of the National Academy of Sciences 96, 5884–5889 (1999).

16. E. E. Noble, C. A. Olson, E. Davis, L. Tsan, Y.-W. Chen, R. Schade, C. Liu, A. Suarez, R. B. Jones, C. De La Serre, X. Yang, E. Y. Hsiao, S. E. Kanoski, Gut microbial taxa elevated by dietary sugar disrupt memory function. Translational Psychiatry 11, (2021).

17. Y. Yang, Z. Zhong, B. Wang, X. Xia, W. Yao, L. Huang, Y. Wang, W. Ding, Early-life high-fat diet-induced obesity programs hippocampal development and cognitive functions via regulation of gut commensal Akkermansia muciniphila. Neuropsychopharmacology 44, 2054–2064 (2019).

18. K. A. Clark, J. M. Alves, S. Jones, A. G. Yunker, S. Luo, R. P. Cabeen, B. Angelo, A. H. Xiang, K. A. Page, Dietary Fructose Intake and Hippocampal Structure and Connectivity during Childhood. Nutrients 12, 909 (2020).

19. C. L. Baym, N. A. Khan, J. M. Monti, L. B. Raine, E. S. Drollette, R. D. Moore, M. R. Scudder, A. F. Kramer, C. H. Hillman, N. J. Cohen, Dietary lipids are differentially associated with hippocampal-dependent relational memory in prepubescent children1,2,3,4. The American Journal of Clinical Nutrition 99, 1026–1033 (2014).

20. A. Ferreira, J. P. Castro, J. P. Andrade, M. Dulce Madeira, A. Cardoso, Cafeteria-diet effects on cognitive functions, anxiety, fear response and neurogenesis in the juvenile rat. Neurobiology of Learning and Memory 155, 197–207 (2018).

21. T. M. Hsu, V. R. Konanur, L. Taing, R. Usui, B. D. Kayser, M. I. Goran, S. E. Kanoski, Effects of sucrose and high fructose corn syrup consumption on spatial memory function and hippocampal neuroinflammation in adolescent rats. Hippocampus 25, 227–239 (2015).

22. S. E. Kanoski, Cognitive and neuronal systems underlying obesity. Physiol Behav 106, 337–344 (2012).

23. S. E. Kanoski, T. L. Davidson, Western diet consumption and cognitive impairment: links to hippocampal dysfunction and obesity. Physiol Behav 103, 59–68 (2011).

24. V. A. Moser, C. J. Pike, Obesity Accelerates Alzheimer-Related Pathology in APOE4 but not APOE3 Mice. eNeuro 4, (2017).

25. R. Molteni, A. Wu, S. Vaynman, Z. Ying, R. J. Barnard, F. Gomez-Pinilla, Exercise reverses the harmful effects of consumption of a high-fat diet on synaptic and behavioral plasticity associated to the action of brain-derived neurotrophic factor. Neuroscience 123, 429–440 (2004).

26. M. E. Hasselmo, The role of acetylcholine in learning and memory. Current Opinion in Neurobiology 16, 710–715 (2006).

27. E. D. Levin, F. J. McClernon, A. H. Rezvani, Nicotinic effects on cognitive function: behavioral characterization, pharmacological specification, and anatomic localization. Psychopharmacology 184, 523–539 (2006).

28. J. G. Bunce, H. R. Sabolek, J. J. Chrobak, Intraseptal infusion of the cholinergic agonist carbachol impairs delayed-non-match-to-sample radial arm maze performance in the rat. Hippocampus 14, 450–459 (2004).

29. F. Fadda, F. Melis, R. Stancampiano, Increased hippocampal acetylcholine release during a working memory task. European Journal of Pharmacology 307, R1–R2 (1996).

30. J. Haam, J. L. Yakel, Cholinergic modulation of the hippocampal region and memory function. Journal of Neurochemistry 142, 111–121 (2017).

31. K.-G. Ma, Y.-H. Qian, Alpha 7 nicotinic acetylcholine receptor and its effects on Alzheimer’s disease. Neuropeptides 73, 96–106 (2019).

32. W. Liu, J. Li, M. Yang, X. Ke, Y. Dai, H. Lin, S. Wang, L. Chen, J. Tao, Chemical genetic activation of the cholinergic basal forebrain hippocampal circuit rescues memory loss in Alzheimer’s disease. Alzheimer’s Research & Therapy 14, (2022).

33. A. Wieckowska-Gacek, A. Mietelska-Porowska, M. Wydrych, U. Wojda, Western diet as a trigger of Alzheimer’s disease: From metabolic syndrome and systemic inflammation to neuroinflammation and neurodegeneration. Ageing research reviews 70, 101397 (2021).

34. E. E. Noble, T. M. Hsu, R. B. Jones, A. A. Fodor, M. I. Goran, S. E. Kanoski, Early-Life Sugar Consumption Affects the Rat Microbiome Independently of Obesity. The Journal of Nutrition 147, 20–28 (2017).

35. L. Tsan, S. Sun, A. M. R. Hayes, L. Bridi, L. S. Chirala, E. E. Noble, A. A. Fodor, S. E. Kanoski, Early life Western diet-induced memory impairments and gut microbiome changes in female rats are long-lasting despite healthy dietary intervention. Nutritional Neuroscience 25, 2490–2506 (2022).

36. L. A. David, C. F. Maurice, R. N. Carmody, D. B. Gootenberg, J. E. Button, B. E. Wolfe, A. V. Ling, A. S. Devlin, Y. Varma, M. A. Fischbach, Diet rapidly and reproducibly alters the human gut microbiome. Nature 505, 559–563 (2014).

37. C. B. de La Serre, C. L. Ellis, J. Lee, A. L. Hartman, J. C. Rutledge, H. E. Raybould, Propensity to high-fat diet-induced obesity in rats is associated with changes in the gut microbiota and gut inflammation. American Journal of Physiology-Gastrointestinal and Liver Physiology 299, G440–G448 (2010).

38. C. De Filippo, D. Cavalieri, M. Di Paola, M. Ramazzotti, J. B. Poullet, S. Massart, S. Collini, G. Pieraccini, P. Lionetti, Impact of diet in shaping gut microbiota revealed by a comparative study in children from Europe and rural Africa. Proceedings of the National Academy of Sciences 107, 14691–14696 (2010).

39. C. A. Olson, A. J. Iñiguez, G. E. Yang, P. Fang, G. N. Pronovost, K. G. Jameson, T. K. Rendon, J. Paramo, J. T. Barlow, R. F. Ismagilov, E. Y. Hsiao, Alterations in the gut microbiota contribute to cognitive impairment induced by the ketogenic diet and hypoxia. Cell Host & Microbe 29, 1378–1392.e1376 (2021).

40. Y. Guo, X. Zhu, M. Zeng, L. Qi, X. Tang, D. Wang, M. Zhang, Y. Xie, H. Li, X. Yang, D. Chen, A diet high in sugar and fat influences neurotransmitter metabolism and then affects brain function by altering the gut microbiota. Transl Psychiatry 11, 328 (2021).

41. N. J. Broadbent, L. R. Squire, R. E. Clark, Spatial memory, recognition memory, and the hippocampus. Proceedings of the National Academy of Sciences 101, 14515–14520 (2004).

42. R. E. Clark, N. J. Broadbent, L. R. Squire, Hippocampus and remote spatial memory in rats. Hippocampus 15, 260–272 (2005).

43. M. C. Martínez, M. E. Villar, F. Ballarini, H. Viola, Retroactive interference of object-in-context long-term memory: Role of dorsal hippocampus and medial prefrontal cortex. Hippocampus 24, 1482–1492 (2014).

44. I. Balderas, C. J. Rodriguez-Ortiz, P. Salgado-Tonda, J. Chavez-Hurtado, J. L. McGaugh, F. Bermudez-Rattoni, The consolidation of object and context recognition memory involve different regions of the temporal lobe. Learning & Memory 15, 618–624 (2008).

45. M. M. Albasser, E. Amin, M. D. Iordanova, M. W. Brown, J. M. Pearce, J. P. Aggleton, Perirhinal cortex lesions uncover subsidiary systems in the rat for the detection of novel and familiar objects. European Journal of Neuroscience 34, 331–342 (2011).

46. S. J. Cohen, R. W. Stackman Jr, Assessing rodent hippocampal involvement in the novel object recognition task. A review. Behavioural Brain Research 285, 105–117 (2015).

47. V. P. Carlini, M. Ghersi, L. Gabach, H. B. Schiöth, M. F. Pérez, O. A. Ramirez, M. Fiol de Cuneo, S. R. de Barioglio, Hippocampal effects of neuronostatin on memory, anxiety-like behavior and food intake in rats. Neuroscience 197, 145–152 (2011).

48. B. R. Miller, R. Hen, The current state of the neurogenic theory of depression and anxiety. Current Opinion in Neurobiology 30, 51–58 (2015).

49. C. André, A.-L. Dinel, G. Ferreira, S. Layé, N. Castanon, Diet-induced obesity progressively alters cognition, anxiety-like behavior and lipopolysaccharide-induced depressive-like behavior: Focus on brain indoleamine 2,3-dioxygenase activation. *Brain*, Behavior, and Immunity 41, 10–21 (2014).

50. G. L. Dunbar, R. J. Rylett, B. M. Schmidt, R. C. Sinclair, L. R. Williams, Hippocampal choline acetyltransferase activity correlates with spatial learning in aged rats. Brain Research 604, 266–272 (1993).

51. P. M. Nagy, I. Aubert, Overexpression of the vesicular acetylcholine transporter enhances dendritic complexity of adult-born hippocampal neurons and improves acquisition of spatial memory during aging. Neurobiology of Aging 36, 1881–1889 (2015).

52. S. Haider, S. Saleem, T. Perveen, S. Tabassum, Z. Batool, S. Sadir, L. Liaquat, S. Madiha, Age-related learning and memory deficits in rats: role of altered brain neurotransmitters, acetylcholinesterase activity and changes in antioxidant defense system. AGE 36, (2014).

53. G. Pepeu, M. G. Giovannini, Cholinesterase inhibitors and memory. Chemico-Biological Interactions 187, 403–408 (2010).

54. S. Kosari, E. Badoer, J. C. D. Nguyen, A. S. Killcross, T. A. Jenkins, Effect of western and high fat diets on memory and cholinergic measures in the rat. Behavioural Brain Research 235, 98–103 (2012).

55. W. H. Meck, R. A. Smith, C. L. Williams, Pre- and postnatal choline supplementation produces long-term facilitation of spatial memory. Developmental Psychobiology 21, 339–353 (1988).

56. I. Martinelli, S. K. Tayebati, P. Roy, M. V. Micioni Di Bonaventura, M. Moruzzi, C. Cifani, F. Amenta, D. Tomassoni, Obesity-Related Brain Cholinergic System Impairment in High-Fat-Diet-Fed Rats. Nutrients 14, 1243 (2022).

57. A. N. Suarez, T. M. Hsu, C. M. Liu, E. E. Noble, A. M. Cortella, E. M. Nakamoto, J. D. Hahn, G. De Lartigue, S. E. Kanoski, Gut vagal sensory signaling regulates hippocampus function through multi-order pathways. Nature Communications 9, (2018).

58. M. Covasa, J. Grahn, R. C. Ritter, High fat maintenance diet attenuates hindbrain neuronal response to CCK. Regul Pept 86, 83–88 (2000).

59. D. M. Savastano, M. Covasa, Adaptation to a high-fat diet leads to hyperphagia and diminished sensitivity to cholecystokinin in rats. J Nutr 135, 1953–1959 (2005).

60. G. de Lartigue, C. Barbier de la Serre, E. Espero, J. Lee, H. E. Raybould, Leptin resistance in vagal afferent neurons inhibits cholecystokinin signaling and satiation in diet induced obese rats. PLoS One 7, e32967 (2012).

61. N. T. Baxter, A. W. Schmidt, A. Venkataraman, K. S. Kim, C. Waldron, T. M. Schmidt, Dynamics of human gut microbiota and short-chain fatty acids in response to dietary interventions with three fermentable fibers. MBio 10, e02566–02518 (2019).

62. F. Fava, R. Gitau, B. A. Griffin, G. Gibson, K. Tuohy, J. Lovegrove, The type and quantity of dietary fat and carbohydrate alter faecal microbiome and short-chain fatty acid excretion in a metabolic syndrome ‘at-risk’population. International journal of obesity 37, 216–223 (2013).

63. J. Tap, J. P. Furet, M. Bensaada, C. Philippe, H. Roth, S. Rabot, O. Lakhdari, V. Lombard, B. Henrissat, G. Corthier, Gut microbiota richness promotes its stability upon increased dietary fibre intake in healthy adults. Environmental microbiology 17, 4954–4964 (2015).

64. A. C. Ericsson, C. L. Franklin, The gut microbiome of laboratory mice: considerations and best practices for translational research. Mammalian Genome 32, 239–250 (2021).

65. C. L. Franklin, A. C. Ericsson, Microbiota and reproducibility of rodent models. Lab Animal 46, 114–122 (2017).

66. A. L. Unger, K. Eckstrom, T. L. Jetton, J. Kraft, Colonic bacterial composition is sex-specific in aged CD-1 mice fed diets varying in fat quality. PLOS ONE 14, e0226635 (2019).

67. Z. Zhang, J. E. Hyun, A. Thiesen, H. Park, N. Hotte, H. Watanabe, T. Higashiyama, K. L. Madsen, Sex-Specific Differences in the Gut Microbiome in Response to Dietary Fiber Supplementation in IL-10-Deficient Mice. Nutrients 12, 2088 (2020).

68. E. E. Noble, T. M. Hsu, J. Liang, S. E. Kanoski, Early-life sugar consumption has long-term negative effects on memory function in male rats. Nutritional Neuroscience 22, 273–283 (2019).

69. M. M. Kaczmarczyk, A. S. Machaj, G. S. Chiu, M. A. Lawson, S. J. Gainey, J. M. York, D. D. Meling, S. A. Martin, K. A. Kwakwa, A. F. Newman, J. A. Woods, K. W. Kelley, Y. Wang, M. J. Miller, G. G. Freund, Methylphenidate prevents high-fat diet (HFD)-induced learning/memory impairment in juvenile mice. Psychoneuroendocrinology 38, 1553–1564 (2013).

70. S. E. Kanoski, T. L. Davidson, Different patterns of memory impairments accompany short- and longer-term maintenance on a high-energy diet. J Exp Psychol Anim Behav Process 36, 313–319 (2010).

71. P. M. Sullivan, Influence of Western diet and APOE genotype on Alzheimer’s disease risk. Neurobiology of disease 138, 104790 (2020).

72. R. Quinn, Comparing rat’s to human’s age: how old is my rat in people years? Nutrition 21, 775 (2005).

73. P. Sengupta, The Laboratory Rat: Relating Its Age With Human’s. Int J Prev Med 4, 624–630 (2013).

74. L. Tsan, S. Chometton, A. M. R. Hayes, M. E. Klug, Y. Zuo, S. Sun, L. Bridi, R. Lan, A. A. Fodor, E. E. Noble, X. Yang, S. E. Kanoski, L. A. Schier, Early life low-calorie sweetener consumption disrupts glucose regulation, sugar-motivated behavior, and memory function in rats. JCI Insight, (2022).

75. E. A. Davis, H. S. Wald, A. N. Suarez, J. Zubcevic, C. M. Liu, A. M. Cortella, A. K. Kamitakahara, J. W. Polson, M. Arnold, H. J. Grill, G. De Lartigue, S. E. Kanoski, Ghrelin Signaling Affects Feeding Behavior, Metabolism, and Memory through the Vagus Nerve. Current Biology 30, 4510–4518.e4516 (2020).

76. L. Décarie-Spain, C. M. Liu, L. T. Lauer, K. Subramanian, A. G. Bashaw, M. E. Klug, I. H. Gianatiempo, A. N. Suarez, E. E. Noble, K. N. Donohue, A. M. Cortella, J. D. Hahn, E. A. Davis, S. E. Kanoski, Ventral hippocampus-lateral septum circuitry promotes foraging-related memory. Cell Reports 40, 111402 (2022).

77. K. S. Subramanian, L. T. Lauer, A. M. R. Hayes, L. Décarie-Spain, K. McBurnett, A. C. Nourbash, K. N. Donohue, A. E. Kao, A. G. Bashaw, D. Burdakov, E. E. Noble, L. A. Schier, S. E. Kanoski, Hypothalamic melanin-concentrating hormone neurons integrate food-motivated appetitive and consummatory processes in rats. Nature Communications 14, (2023).

78. S. B. Caine, M. A. Geyer, N. R. Swerdlow, Carbachol infusion into the dentate gyrus disrupts sensorimotor gating of startle in the rat. Psychopharmacology 105, 347–354 (1991).

## Supplementary References

1. E. A. Davis, H. S. Wald, A. N. Suarez, J. Zubcevic, C. M. Liu, A. M. Cortella, A. K. Kamitakahara, J. W. Polson, M. Arnold, H. J. Grill, G. De Lartigue, S. E. Kanoski, Ghrelin Signaling Affects Feeding Behavior, Metabolism, and Memory through the Vagus Nerve. Current Biology 30, 4510–4518.e4516 (2020).

2. L. Tsan, S. Sun, A. M. R. Hayes, L. Bridi, L. S. Chirala, E. E. Noble, A. A. Fodor, S. E. Kanoski, Early life Western diet-induced memory impairments and gut microbiome changes in female rats are long-lasting despite healthy dietary intervention. Nutritional Neuroscience 25, 2490–2506 (2022).

3. L. Tsan, S. Chometton, A. M. R. Hayes, M. E. Klug, Y. Zuo, S. Sun, L. Bridi, R. Lan, A. A. Fodor, E. E. Noble, X. Yang, S. E. Kanoski, L. A. Schier, Early life low-calorie sweetener consumption disrupts glucose regulation, sugar-motivated behavior, and memory function in rats. JCI Insight, (2022).

4. A. M. R. Hayes, L. Tsan, A. E. Kao, G. M. Schwartz, L. Décarie-Spain, L. Tierno Lauer, M. E. Klug, L. A. Schier, S. E. Kanoski, Early Life Low-Calorie Sweetener Consumption Impacts Energy Balance during Adulthood. Nutrients 14, 4709 (2022).

5. M. M. Albasser, E. Amin, M. D. Iordanova, M. W. Brown, J. M. Pearce, J. P. Aggleton, Perirhinal cortex lesions uncover subsidiary systems in the rat for the detection of novel and familiar objects. European Journal of Neuroscience 34, 331–342 (2011).

6. E. E. Noble, C. A. Olson, E. Davis, L. Tsan, Y.-W. Chen, R. Schade, C. Liu, A. Suarez, R. B. Jones, C. De La Serre, X. Yang, E. Y. Hsiao, S. E. Kanoski, Gut microbial taxa elevated by dietary sugar disrupt memory function. Translational Psychiatry 11, (2021).

7. J. E. Beilharz, J. Maniam, M. J. Morris, Short exposure to a diet rich in both fat and sugar or sugar alone impairs place, but not object recognition memory in rats. *Brain*, Behavior, and Immunity 37, 134–141 (2014).

8. A. N. Suarez, T. M. Hsu, C. M. Liu, E. E. Noble, A. M. Cortella, E. M. Nakamoto, J. D. Hahn, G. De Lartigue, S. E. Kanoski, Gut vagal sensory signaling regulates hippocampus function through multi-order pathways. Nature Communications 9, (2018).

9. C. Belzung, G. Griebel, Measuring normal and pathological anxiety-like behaviour in mice: a review. Behavioural Brain Research 125, 141–149 (2001).

10. J. Chen, K. E. Cho, D. Skwarzynska, S. Clancy, N. J. Conley, S. M. Clinton, X. Li, L. Lin, J. J. Zhu, The Property-Based Practical Applications and Solutions of Genetically Encoded Acetylcholine and Monoamine Sensors. The Journal of Neuroscience 41, 2318–2328 (2021).

11. L. Lin, S. Gupta, W. S. Zheng, K. Si, J. J. Zhu, Genetically encoded sensors enable micro- and nano-scopic decoding of transmission in healthy and diseased brains. Molecular Psychiatry 26, 443–455 (2021).

12. P. K. Zhu, W. S. Zheng, P. Zhang, M. Jing, P. M. Borden, F. Ali, K. Guo, J. Feng, J. S. Marvin, Y. Wang, J. Wan, L. Gan, A. C. Kwan, L. Lin, L. L. Looger, Y. Li, Y. Zhang, Nanoscopic Visualization of Restricted Nonvolume Cholinergic and Monoaminergic Transmission with Genetically Encoded Sensors. Nano Letters 20, 4073–4083 (2020).

13. P. M. Borden, P. Zhang, A. V. Shivange, J. S. Marvin, J. Cichon, C. Dan, K. Podgorski, A. Figueiredo, O. Novak, M. Tanimoto, E. Shigetomi, M. A. Lobas, H. Kim, P. K. Zhu, Y. Zhang, W. S. Zheng, C. Fan, G. Wang, B. Xiang, L. Gan, G.-X. Zhang, K. Guo, L. Lin, Y. Cai, A. G. Yee, A. Aggarwal, C. P. Ford, D. C. Rees, D. Dietrich, B. S. Khakh, J. S. Dittman, W.-B. Gan, M. Koyama, V. Jayaraman, J. F. Cheer, H. A. Lester, J. J. Zhu, L. L. Looger. (Cold Spring Harbor Laboratory, 2020).

14. A. L. Nichols, Z. Blumenfeld, C. Fan, L. Luebbert, A. E. M. Blom, B. N. Cohen, J. S. Marvin, P. M. Borden, C. H. Kim, A. K. Muthusamy, A. V. Shivange, H. J. Knox, H. R. Campello, J. H. Wang, D. A. Dougherty, L. L. Looger, T. Gallagher, D. C. Rees, H. A. Lester, Fluorescence activation mechanism and imaging of drug permeation with new sensors for smoking-cessation ligands. eLife 11, e74648 (2022).

15. L. W. Swanson, Brain maps 4.0-Structure of the rat brain: An open access atlas with global nervous system nomenclature ontology and flatmaps. J Comp Neurol 526, 935–943 (2018).

16. S. E. Kanoski, S. M. Fortin, K. M. Ricks, H. J. Grill, Ghrelin Signaling in the Ventral Hippocampus Stimulates Learned and Motivational Aspects of Feeding via PI3K-Akt Signaling. Biological Psychiatry 73, 915–923 (2013).

17. T. M. Hsu, V. R. Konanur, L. Taing, R. Usui, B. D. Kayser, M. I. Goran, S. E. Kanoski, Effects of sucrose and high fructose corn syrup consumption on spatial memory function and hippocampal neuroinflammation in adolescent rats. Hippocampus 25, 227–239 (2015).

18. J. G. Caporaso, C. L. Lauber, W. A. Walters, D. Berg-Lyons, C. A. Lozupone, P. J. Turnbaugh, N. Fierer, R. Knight, Global patterns of 16S rRNA diversity at a depth of millions of sequences per sample. Proceedings of the National Academy of Sciences 108, 4516–4522 (2011).

19. E. Bolyen, J. R. Rideout, M. R. Dillon, N. A. Bokulich, C. C. Abnet, G. A. Al-Ghalith, H. Alexander, E. J. Alm, M. Arumugam, F. Asnicar, Y. Bai, J. E. Bisanz, K. Bittinger, A. Brejnrod, C. J. Brislawn, C. T. Brown, B. J. Callahan, A. M. Caraballo-Rodríguez, J. Chase, E. K. Cope, R. Da Silva, C. Diener, P. C. Dorrestein, G. M. Douglas, D. M. Durall, C. Duvallet, C. F. Edwardson, M. Ernst, M. Estaki, J. Fouquier, J. M. Gauglitz, S. M. Gibbons, D. L. Gibson, A. Gonzalez, K. Gorlick, J. Guo, B. Hillmann, S. Holmes, H. Holste, C. Huttenhower, G. A. Huttley, S. Janssen, A. K. Jarmusch, L. Jiang, B. D. Kaehler, K. B. Kang, C. R. Keefe, P. Keim, S. T. Kelley, D. Knights, I. Koester, T. Kosciolek, J. Kreps, M. G. I. Langille, J. Lee, R. Ley, Y.-X. Liu, E. Loftfield, C. Lozupone, M. Maher, C. Marotz, B. D. Martin, D. McDonald, L. J. McIver, A. V. Melnik, J. L. Metcalf, S. C. Morgan, J. T. Morton, A. T. Naimey, J. A. Navas-Molina, L. F. Nothias, S. B. Orchanian, T. Pearson, S. L. Peoples, D. Petras, M. L. Preuss, E. Pruesse, L. B. Rasmussen, A. Rivers, M. S. Robeson, P. Rosenthal, N. Segata, M. Shaffer, A. Shiffer, R. Sinha, S. J. Song, J. R. Spear, A. D. Swafford, L. R. Thompson, P. J. Torres, P. Trinh, A. Tripathi, P. J. Turnbaugh, S. Ul-Hasan, J. J. J. Van Der Hooft, F. Vargas, Y. Vázquez-Baeza, E. Vogtmann, M. Von Hippel, W. Walters, Y. Wan, M. Wang, J. Warren, K. C. Weber, C. H. D. Williamson, A. D. Willis, Z. Z. Xu, J. R. Zaneveld, Y. Zhang, Q. Zhu, R. Knight, J. G. Caporaso, Reproducible, interactive, scalable and extensible microbiome data science using QIIME 2. Nature Biotechnology 37, 852–857 (2019).

20. B. J. Callahan, P. J. McMurdie, M. J. Rosen, A. W. Han, A. J. A. Johnson, S. P. Holmes, DADA2: High-resolution sample inference from Illumina amplicon data. Nature Methods 13, 581–583 (2016).

21. R. B. Jones, X. Zhu, E. Moan, H. J. Murff, R. M. Ness, D. L. Seidner, S. Sun, C. Yu, Q. Dai, A. A. Fodor, M. A. Azcarate-Peril, M. J. Shrubsole, Inter-niche and inter-individual variation in gut microbial community assessment using stool, rectal swab, and mucosal samples. Scientific Reports 8, (2018)

